# Exogenous DCPTA ameliorates the soil drought effect on nitrogen metabolism in maize during the pre-female inflorescence emergence stage

**DOI:** 10.1101/385203

**Authors:** Tenglong Xie, Wanrong Gu, Zhang Liguo, Congfeng Li, Wenhua Li, Shi Wei

## Abstract

**Abstract:** 2-(3,4-Dichlorophenoxy) triethylamine (DCPTA) regulates many aspects of plant development; however, its effects on soil drought tolerance are unknown. We pre-treated maize (*Zea mays* L.) by foliar application of DCPTA and subsequently exposed the plants to soil drought and rewatering conditions during the pre-female inflorescence emergence stage. Exogenous DCPTA significantly alleviated drought-induced decreases in maize yield, shoot and root relative growth rate (RGR), leaf relative water content (RLWC), net photosynthetic rate (Pn), stomatal conductance (Gs) and transpiration rate (Tr), nitrate (NO_3_^−^), nitrite (NO_2_^−^), and soluble protein contents, and nitrate reductase (NR), nitrite reductase (NiR), isocitrate dehydrogenase (ICDH), alanine aminotransferase (AlaAT) and aspartate aminotransferase (AspAT) activities; increases in the intercellular CO_2_ concentration (Ci), the ammonium (NH_4_^+^) and free amino acid contents, and the glutamate dehydrogenase (GDH) and protease activities. Simultaneously, exogenous DCPTA improved the spatial and temporal distribution of roots and increased the root hydraulic conductivity (Lp), flow rate of root-bleeding sap and NO_3_^−^ delivery rates. Moreover, Exogenous DCPTA protected the chloroplast structure from drought injury. Taken together, our results suggest that exogenous DCPTA mitigates the repressive effects of drought on N metabolism and subsequently enhances drought tolerance during the pre-female inflorescence emergence stage of maize.

**Highlights:** This is the first article that explores the effects of DCPTA on nitrogen metabolism and the first article that explores the effects of DCPTA on crops under soil drought conditions.

## Introduction

Crops are frequently exposed to drought during the growth period because of limited and erratic rainfall patterns due to global climate change, which leads to restrictions on agricultural productivity worldwide (Trenberth *et al.,* 2014). Maize (*Zea mays* L.), an essential component of global food security, is widely cultivated around the word. The majority of the cultivated area of maize is almost wholly rain-fed and experiences sporadic drought and rewetting cycles (Nuccio *et al.,* 2015). However, maize is considered to be a drought-sensitive crop and loses approximately 1/4 potential yield annually due to drought (Ziyomo and Bernardo, 2013). By 2050, the world population will reach 9 billion people, resulting in a high demand for maize (projected to double); furthermore, at that time, drought will severely restrict crop growth for more than 50% of the cultivated land (Schiermeier, 2014).

To stabilize and increase global crop production to satisfy the demand of the globally burgeoning population, it is imperative to design agronomic research to improve maize performance under drought stress (Zhao *et al.,* 2018). The application of plant growth regulators has been considered an effective way to enhance crop drought resistance (Ali *et al.,* 2017). Multiple investigations have indicated that a tertiary amine bioregulator known as 2-(3,4-dichlorophenoxy) triethylamine (DCPTA) regulates many aspects of plant development; for example, DCPTA promotes plant growth (Keithly *et al.,* 1990a), enlarges chloroplast volume (Keithly *et al.,* 1990b), enhances photosynthetic enzyme activity (Wan and Mendoza, 1992), accelerates CO_2_ fixation (Gausman *et al.,* 1985), and stimulates carotenoid biosynthesis (Benedict *et al.,* 1985). As far as we know, very few studies of DCPTA have focused on crops, and the effect of DCPTA on crops exposed to soil drought are still unclear.

Nitrogen (N) metabolism is a fundamental process in determining the growth and productivity of plants (Kusano *et al.,* 2011). After being taken up by root systems, nitrate (NO_3_^−^) is converted to nitrite (NO_2_^−^) by nitrate reductase (NR), the first step of N uptake and utilization. Subsequently, nitrite (NO_2_^−^) is converted to NH_4_+ by nitrite reductase (NiR) with reduction-ferredoxin (Fd_red_) as an electron donor (Rajasekhar *et al.*, 2010). Afterward, the ammonium (NH_4_^+^), derived from NO_3_^−^ reduction, photorespiration and/or other metabolic processes is assimilated into glutamine by the glutamine synthase/glutamine oxoglutarate aminotransferase (GS/GOGAT) cycle or the alternative glutamate dehydrogenase (GDH) pathway with 2-oxoglutarate (2-OG) and reducing equivalents provided by photosynthesis (Chardon *et al*., 2012). Subsequently, glutamate serving as a donor of the amino group is used for the synthesis of other amino acids, which are used for the synthesis of various organic molecules such as chlorophyll, proteins and nucleic acids. The reactions are catalysed by aminotransferases such as alanine aminotransferase (AlaAT) and aspartate aminotransferase (AspAT) (Slattery *et al*., 2017).

Drought disrupts N metabolism mainly via inhibiting the uptake and/or long-distance transportation of NO_3_^−^ (Nacer *et al*., 2013), altering the activities of enzymes involved in N metabolism (Robredo *et al.,* 2011), inhibiting amino acid synthesis, and promoting protein hydrolysis (Fresneau *et al.,* 2007). At present, the study of plant growth regulators mainly concentrates on the improvement of photosynthesis and antioxidant systems, and there have been only a limited number of publications related to N metabolism.

Our previous hydroponic trial found that exogenous DCPTA drastically promoted growth under non-stress conditions and mitigated the PEG-simulated drought-induced growth inhibition of maize at the seedling stage (Xie *et al.,* 2017). The present study was conducted to explore whether DCPTA can alleviate soil drought injuries to maize and whether the effects are associated with the modulation of nitrogen metabolism.

## Materials and Methods

### Plant material, growth conditions, design and sampling

Seeds of the maize cultivar ZhengDan 958 and DCPTA were obtained from the Henan Academy of Agricultural Sciences in China and the China Zhengzhou Zhengshi Chemical Limited Company, respectively.

These experiments were performed in 2016 and 2017 at the Experimental Station of Northeast Agricultural University, Harbin (126°73’E, 45°73’N), Heilongjiang province, China. The research field area has a temperate continental monsoon climate. The rainfall and mean temperature data during the study period (2016 and 2017, May–October) are listed in Fig. 1. Pits (inner length, 10 m; width, 7 m; and height, 1.2 m) in the field were used as experiment containers (Fig. 2). Plastic sheets were used to cover the inner sides of the pits, and a rain-proof shed was used to ensure the crops were solely dependent on soil moisture and irrigation over the course of the experiment to maintain the soil water conditions. The soil used was Chernozem and was sieved (pore size, 1 cm) and diluted with vermiculite (particle diameter, 4-8 mm; soil to vermiculite, v/v, 2:1). Before planting, soil chemical analysis was conducted according to Cottenie *et al*. (1982), and the results are presented in Table 1. Fertilization was carried out by adding ammonium nitrate (33.5% N), calcium superphosphate (15.5% P2O5), and potassium sulfate (48% K2O) at the rates of 8.0, 8.0 and 20 kg pit^−1^, respectively, before planting. No fertilizer was applied after planting. All containers were watered to 85% before planting. The seeds were manually sown on 2nd May 2016 and 4th May 2017 and were harvested on 7th October 2016 and 3rd October 2017, respectively. Three seeds were sowed per hole to ensure germination, and only the healthiest seedling within 20 days was kept at each site. Each container consisted of 10 rows, and the plant-to-plant and row-to-row distances were 20 cm and 65 cm, respectively. In addition, the ground around the containers was manually sown with the same plant-to-plant and row-to-row distances. The control of plant diseases and insect pests was conducted by managers. The maize at the nine-leaf stage (during the pre-female inflorescence emergence stage) were treated as follows:

1. plants were irrigated continuously and sprayed with either 10 mL water (well-watered) or DCPTA (well-watered+DCPTA) per plant;
2. irrigation was stopped to form the drought conditions and sprayed with 10 mL of either water (drought) or DCPTA (drought +DCPTA) per plant; plants were rehydrated after 20 days of drought treatment.

**Table 1.** Chemical properties of the used soil.

| Year | pH | HCO<sub>3</sub><sup>-</sup><br>+<br>CO<sub>3</sub><sup>2-</sup><br>(mg<br>kg<sup>-1</sup>) | Cl<sup>-</sup><br>(mg<br>kg<sup>-1</sup>) | SO<sub>4</sub><sup>2-</sup><br>(mg<br>kg<sup>-1</sup>) | Ca<sub>2</sub><sup>+</sup><br>(mg<br>kg<sup>-1</sup>) | Mg<sub>2</sub><sup>+</sup><br>(mg<br>kg<sup>-1</sup>) | Na<sup>+</sup><br>(mg<br>kg<sup>-1</sup>) | K<sup>+</sup><br>(mg<br>kg<sup>-1</sup>) | N<br>(mg<br>kg<sup>-1</sup>) | P<br>(mg<br>kg<sup>-1</sup>) |
| --- | --- | --- | --- | --- | --- | --- | --- | --- | --- | --- |
| 2016 | 7.2 | 204.3 | 297.6 | 463.8 | 87.4 | 37.6 | 4.1 | 28.9 | 17.3 | 3.9 |
| 2017 | 7.1 | 201.7 | 302.5 | 446.2 | 90.3 | 40.9 | 4.0 | 31.4 | 16.1 | 3.8 |

**Fig. 1.**
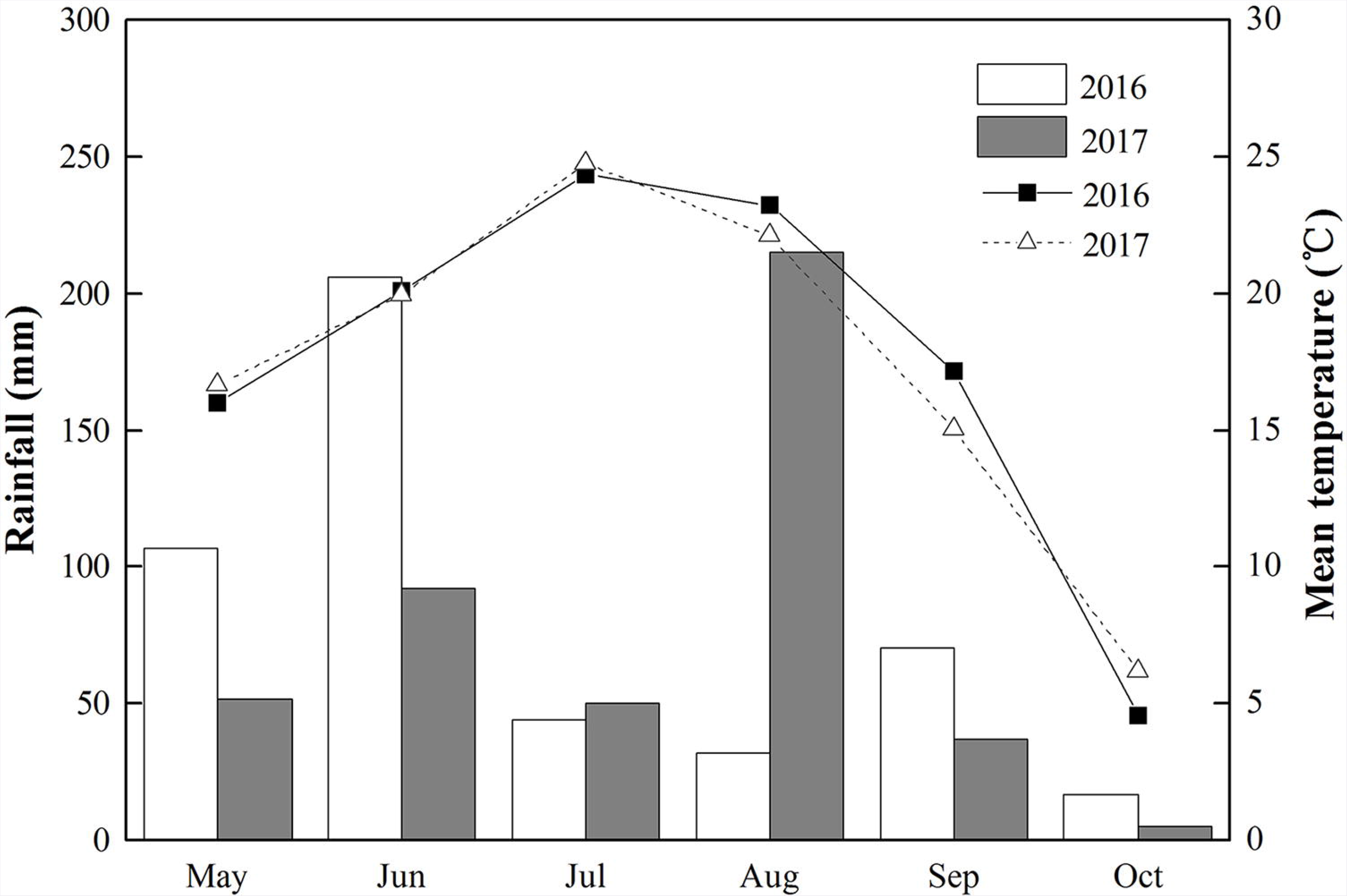
The rainfall (bar) and mean temperature (line) data during the study period (2016 and 2017, May–October).

**Fig. 2.**
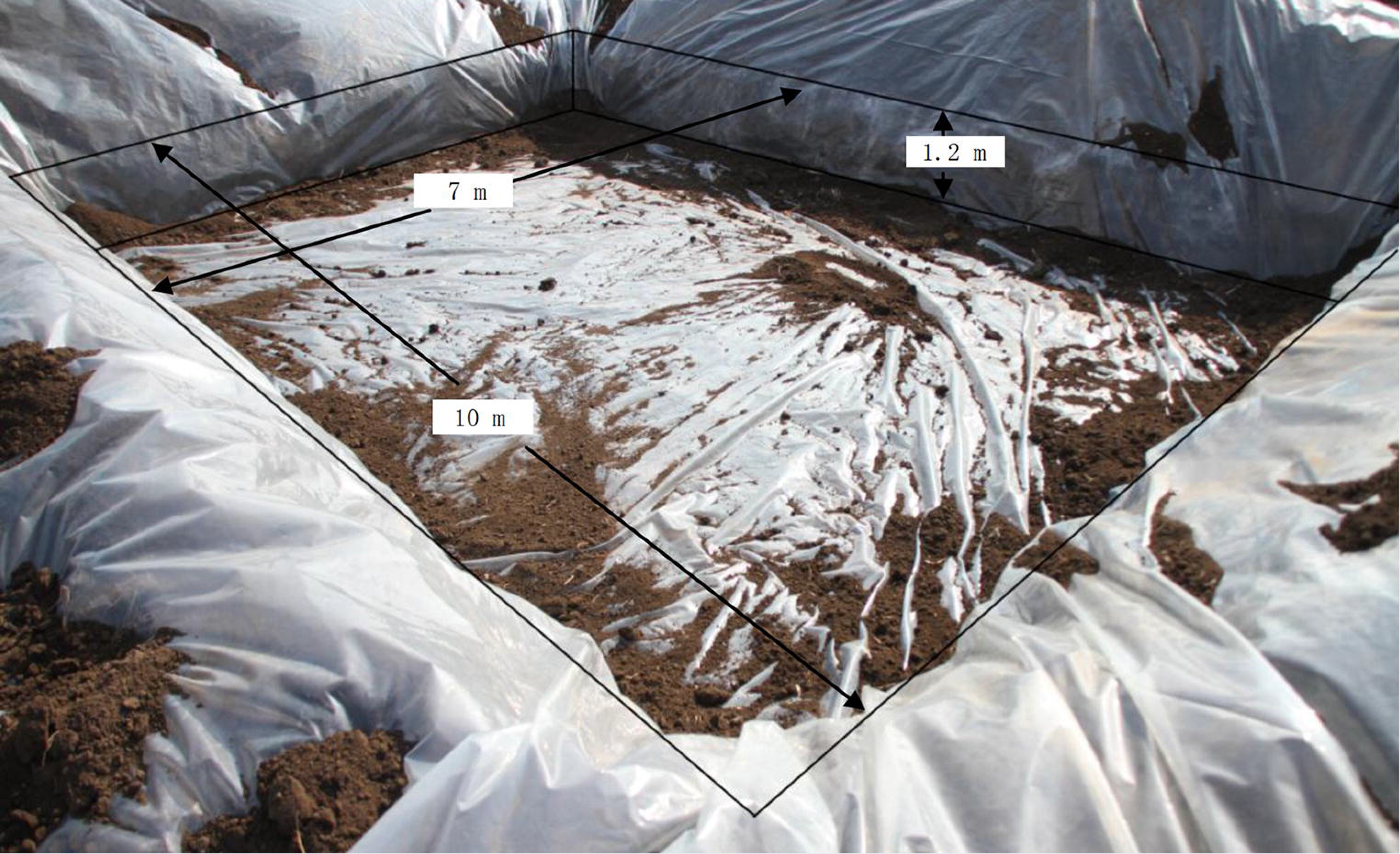
The pits (inner length, 10 m; width, 7 m; and height, 1.2 m) used for this study and the plastic sheets used to cover inner sides of the pits.

The concentrations of DCPTA (25 mg/L) were based on the results of previous screening experiments, and Tween-20 (0.03%) was added as a surfactant to the solution for spraying. Each treatment had five replicates, and experiments were performed in a completely randomized design. The dynamic changes in soil water contents during the experimental stage are exhibited in Fig. 3.

**Fig. 3.**
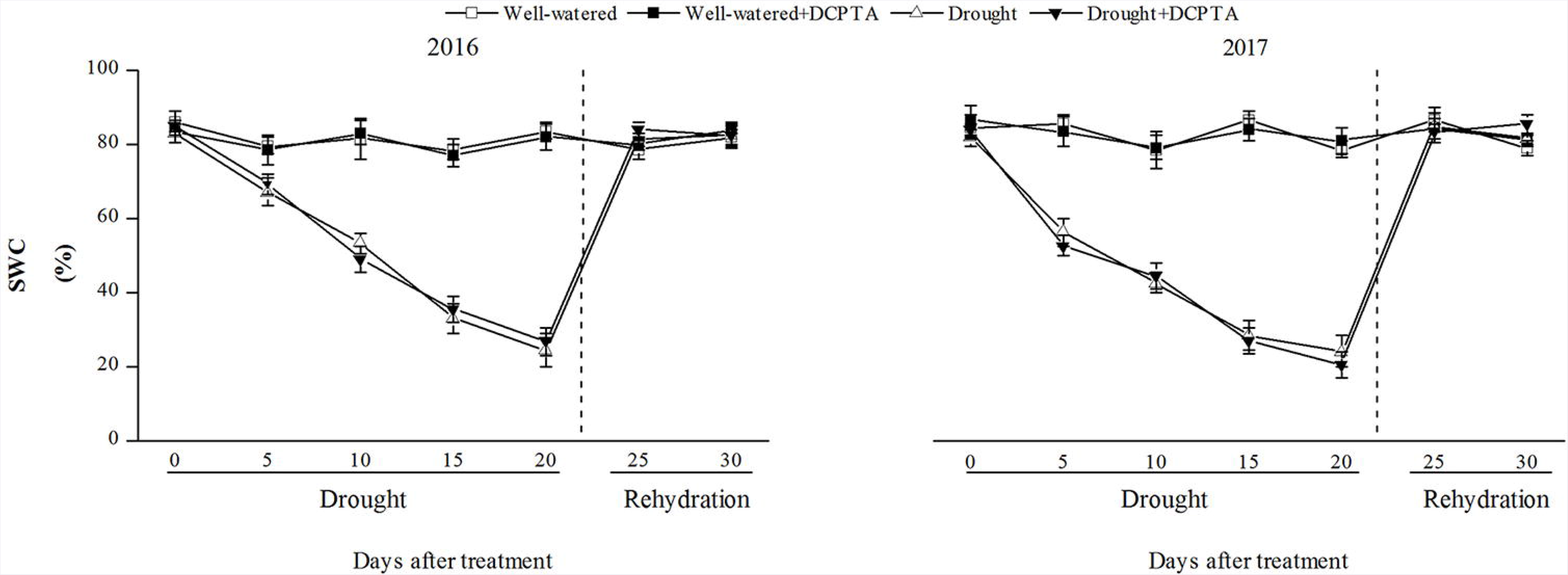
Changes in the soil water content (SWC) in 2016 and 2017. The data represent the means of independent measurements with five replicates, and the standard deviations are indicated by the vertical error bars. Values with the same letters on the bars are not significantly different at P<0.05 (LSD test).

Random plants from each treatment were sampled on days 0, 5, 10, 15, 20, 25 and 30. For leaf sampling, the middle part of the 9^th^ leaf (numbered basipetally) was sampled for analysis of leaf gas exchange, and the same part of the leaf was stored at-80°C after immersion in liquid nitrogen for 30 min for determination of physiological parameters. For root sampling, a hand-held soil auger (inner diameter of 20 cm) was used to obtain soil cores from 0 to100 cm depth of the soil profile at 10 cm increments. The soil cores were soaked in a plastic container overnight, and roots were stirred and sieved through a mesh (400 holes cm^−2^). The soil cores were the carefully washed by swirling water through the cores. The soil material and old dead roots debris were manually separated from the live roots.

## Plant measurement and analysis

### Relative growth rate (RGR) and plant productivity

The shoots and roots of maize were oven dried at 105°C for 45 min and then held at 80°C for 48 h; the shoot and root dry weights plant^−1^ were determined soon afterwards. The RGR was determined as follows: RGR (fresh weight) = [ln (final dry weight) – ln (initial dry weight)]/(duration of treatment days) (Kingsbury *et al.,* 1984). A leaf area metre (Li-COR 3100; Li-COR, Lincoln, NE, USA) was used to estimate the leaf area; number of grains plant^−1^ (GN) and grain yield plant^−1^ (GY) were recorded at the maize physiological maturity stage.

### Leaf relative water content of (RLWC) and soil water content (SWC)

The RLWC was determined on fresh leaf disks (2×2 cm) from the middle part of the 8th leaves (numbered basipetally). After they were weighed (FW), the disks were immersed in distilled water at 25 °C overnight to obtain the turgid weight (TW). The leaves were dried at 80°C for 48 h and then weighed a third time (DW). RLWC was calculated as follows:

RLWC (%) = [(FW-DW)/ (TW - DW)] × 100.

SWC was determined in the soil from the internal area of each container. After being weighed (FW), the soil portion was dried at 85°C for 96 h and then weighed (DW). SWC was calculated as follows:

SWC (%) = [(FW-DW)/DW] ×100.

### Gas exchange

The photosynthetic rate (Pn), transpiration rate (Tr), stomatal conductance (Gs), and intercellular CO2 concentration (Ci) values were determined with a portable photosynthesis system (LI-6400XT; LI-COR Biosciences, Lincoln, NE, USA) between at 13:00~14:00 h. The 6-cm^2^ leaf chamber was used, and the photo flux density was 1000 lmol m^−2^s^−1^.

### Transmission electron microscopy of chloroplasts

Observations were performed according to the description of Hu *et al*. (2014), and the chloroplast ultrastructure was observed under a H-7650 transmission electron microscope (manufacture: Hitachi, Japan).

### Root morphological traits, root hydraulic conductivity, and the collection of root-bleeding sap

Roots from each soil core were scanned using a digital scanner (Epson V700, Indonesia). The root images were analysed using the WinRHIZO Image Analysis system (Version 2013e) (Regent Instruments Inc., Canada). The root length density _32 —3 (RLD, cm root cm^−3^ soil) and root square area density (RSD, cm root cm soil) were calculated according to the method described by Mosaddeghi *et al*. (2009).

The hydrostatic root hydraulic conductivity (Lp) was measured with a Scholander pressure chamber according to the method described by López-Pérez *et al*. (2007).

The plants were cut by scissors at 10–12 cm above the soil surface at 18:00-19:00. Centrifuge tubes (inner diameter 40 mm) with cotton were placed on the upper end of the stalks, and the stalk joints and centrifuge tubes were wrapped by plastic film to keep impurities and insects out (Fig. 4). The bleeding sap was collected for 12 h; then, the cotton was extracted from each centrifuge tube and placed into a glass syringe (100 ml), and the root-bleeding sap was squeezed out for volume measurement. The NO_3_^−^ content in the root-bleeding sap was determined by AA3 Continuous Flow Analytical System (Seal, Germany) according to Guan *et al*. (2014)’ The flow rate of the root-bleeding sap and the NO_3_^−^ delivery rate were expressed as ml h^−1^ root^−1^ and μg h^—1^ root^−1^, respectively.

**Fig. 4.**
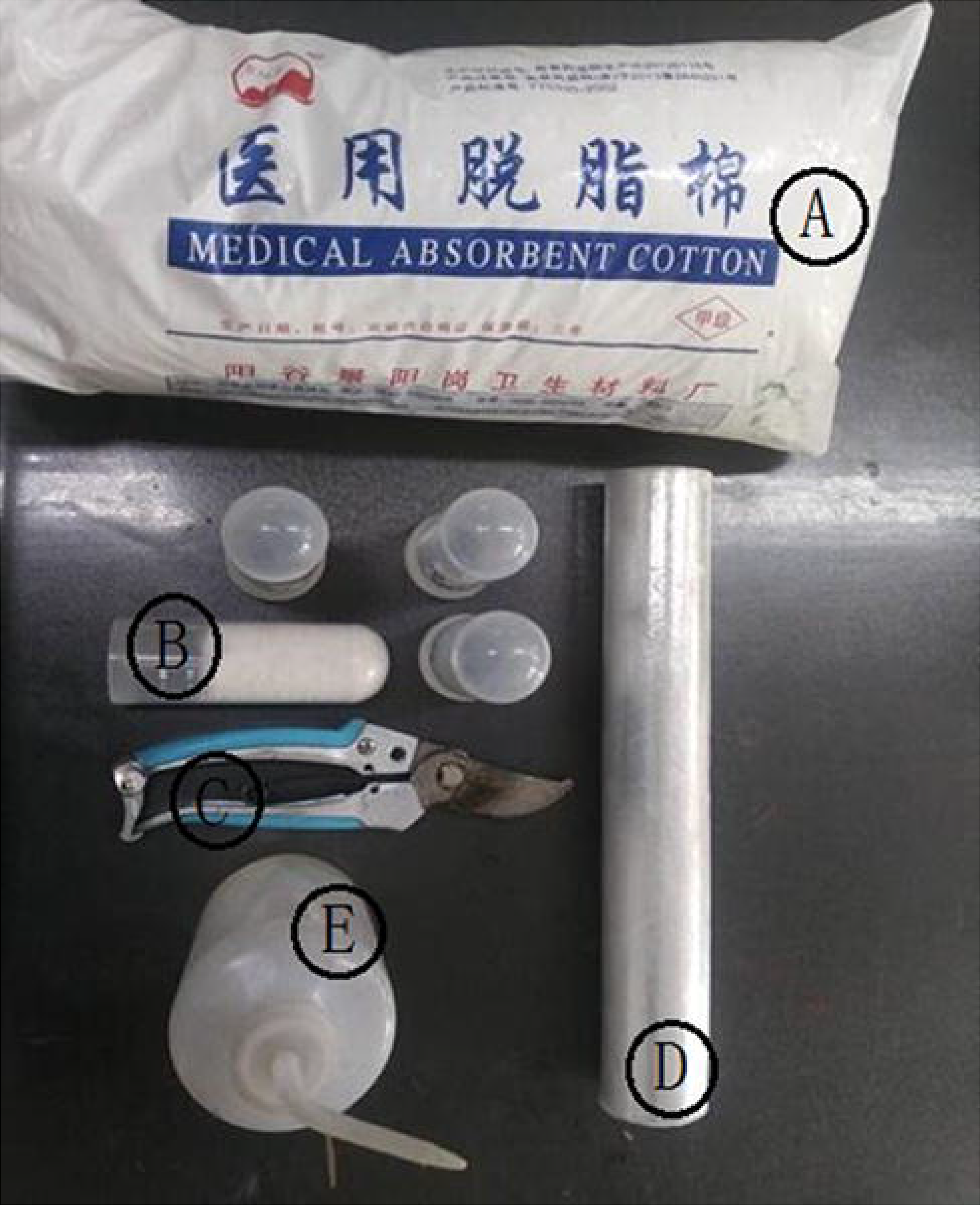
The tools used for the collection of bleeding sap. (A) cotton, (B) centrifuge tube, (C) scissors, (D) plastic film, and (E) deionized water.

### Foliar NO3”, NO_2_^−^” and NH_4_^+^ contents

The foliar NO_3_^−^ content determination by the reduction of NO_3_^−^ to NO_2_^−^ followed the salicylic acid methods of Cataldo *et al*. (1975), the absorbance was monitored at 410 nm. The foliar NO_2_^−^ content was determined using the method described by Barro *et al*. (1991). The NO_2_^−^ content was calculated according to the standard curve obtained by known concentrations of KNO_3_. The NH_4_^+^ content was determined by measuring the absorbance changes at 620 nm, as described by Weber (2007). The NH_4_^+^ content was calculated according to the standard curve obtained by known concentrations of (NH_4_)_2_SO_4_.

### Enzyme activities involved in nitrogen metabolism

The activities of foliar NR and NiR were measured based on the conversion of NO_3_^−^ to NO_2_^−^ and NO_3_^−^ to NH_4_^+^, following the methods of Barro *et al*. (1991), and Ida and Morita (1973), respectively. One NR and NiR unit was expressed as the enzyme amount required for the conversion of 1 mmol of NO_3_^−^ to NO_2_^−^ per hour and the enzyme amount required for the conversion of 1 mmol of NO_3_^−^ to NH_4_^+^ min^−1^, respectively. GS activity was measured according to the methods of O’Neal and Joy (1973). One unit of GS was expressed as the enzyme amount required to catalyse the formation of 1 mmol of glutamylmonohydroxamate per min. The results were expressed as μmol NADH used per minute per milligram of Pro. The GOGAT activity was measured based on the conversion of 2-ketoglutarate to glutamate, following the methods of Groat and Vance (1981). One unit of GOGAT was expressed as the number of enzymes catalysing the oxidation of 1 mmol of NADH per min. The deaminating GDH activity (NAD-GDH) and aminating GDH activity (NADH-GDH) were determined by recording the reduction of NAD and the oxidation of NADH, respectively, as described by Groat and Vance (1981). One unit of GDH was calculated in units of mmol of NADH oxidized/NAD reduced per minute. Isocitrate dehydrogenase (ICDH) was assayed according to the method of Lòpez-Millàn *et al*. (2000). The activity was expressed as U per minute per milligram of protein.

### Activity of AlaAT and AspAT

The AlaAT and AspAT activities were determined according to the methods of Jia *et al*. (2015). Enzyme activity was expressed as μmol g^−1^min^−1^.

### Contents of free amino acids and soluble proteins and protease activity

The free amino acid contents were assayed with the ninhydrin reagent method according to Yemm and Cocking (1955), followed by absorbance readings at 570 nm using glycine as the standard. The soluble protein contents were determined using the Coomassie Brilliant Blue G-250 reagent following the description of Bradford (1976), followed by absorbance readings at 595 nm using bovine serum albumin as the standard. Protease activity was determined by the casein digestion assay described by Drapeau (1974). By this method, one unit is the number of enzymes required to release acid-soluble fragments equivalent to 0.001 A280 per minute at 37°C and pH 7.8.

### Statistical analysis

The data were analysed using SPSS 17.0 and all the values are presented as the mean ± SE. The means were separated using the least significant difference (LSD) test at the 5% probability level.

## Results

### Yield

Drought stress significantly inhibited the maize yield (Fig. 5). Compared with the well-watered treatment, in the drought treatment, the grain number decreased by 29.23% in 2016 and 33.24% in 2017, and the grain yield decreased by 34.06% in 2016 and by 38.22% in 2017. However, the decrease in maize yield was partially recovered by DCPTA. Compared with the well-watered treatment, in the drought+DCPTA treatment, the grain number decreased by 14.01% in 2016 and by 16.55% in 2017, and the grain yield decreased by 17.98% in 2016 and by 20.54% in 2017. Moreover, the application of DCPTA improved maize yield under well-watered conditions. Compared with the well-watered treatment, in the DCPTA treatment, the grain number increased by 5.97% in 2016 and by 6.50% in 2017, and the grain yield increased by 7.31% in 2016 and by 8.02% in 2017.

**Fig. 5.**
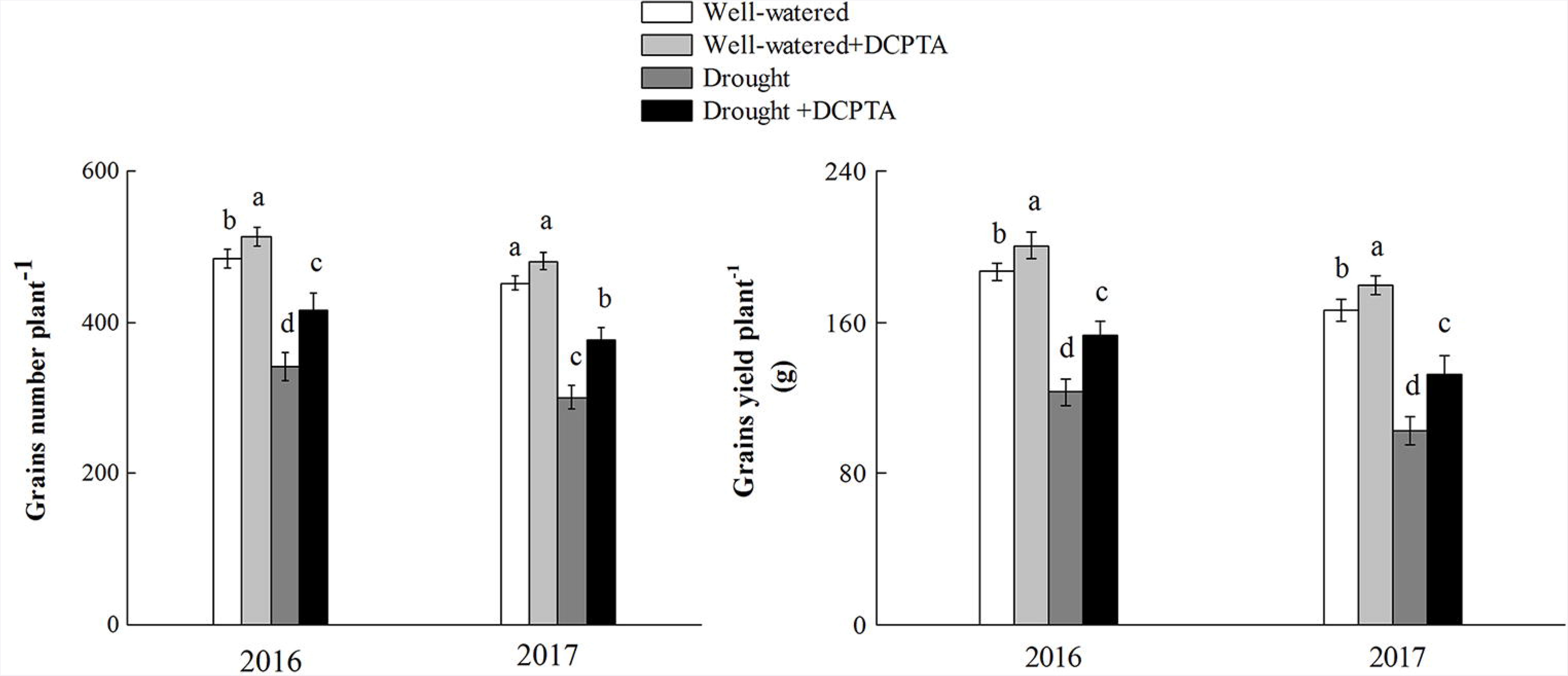
Changes in the grain number plant^−1^ and grain yield plant^−1^ (g) of maize in 2016 and 2017. The data represent the means of independent measurements with five replicates, and the standard deviations are indicated by the vertical error bars. Values with the same letters on the bars are not significantly different at P<0.05 (LSD test).

### Relative growth rate (RGR) of shoot and root

The shoot growth rate was inhibited during the drought period, the root growth rate was promoted over days 0-10 and subsequently decreased, and the growth rates of shoots and roots recovered during rehydration (Fig. 6 and 7). In the drought treatment compared with the control in 2016, the RGR of shoots and roots decreased by 52.26% and by 48.59%, respectively, between day 15 and 20 and by 39.76% and 37.70%, respectively, between day 25 and 30 in 2016. In the drought treatment compared with the control in 2017, the RGR of shoots and roots decreased by 65.18% and 66.52%, respectively, between day 15 and 20 and by 51.97% and 43.74%, respectively, between day 25 and 30 in 2017. However, the decrease in growth was partially recovered by DCPTA. In the drought+DCPTA treatment compared with the control in 2016, the RGR of shoots and roots decreased by 29.96% and 26.31%, respectively, between day 15 and 20, and by 10.11% and 11.49%, respectively, between day 25 and 30. In the drought+DCPTA treatment compared with the control in 2017, the RGR of shoots and roots decreased by 34.20% and 42.23%, respectively, between day 15 and 20, and by 12.41% and 14.75%, respectively, between day 25 and 30. Moreover, the application of DCPTA improved maize growth under well-watered conditions. The shoot RGR difference between the well-watered+DCPTA treatment and well-watered treatment was significant at days 16-20 and 26-30 in 2016 and days 11-30 in 2017, respectively. Similarly, the root RGR difference between the DCPTA treatment and well-watered treatment was significant at days 6-10 and 16-25 in 2016 and days 11-30 in 2017, respectively.

**Fig. 6.**
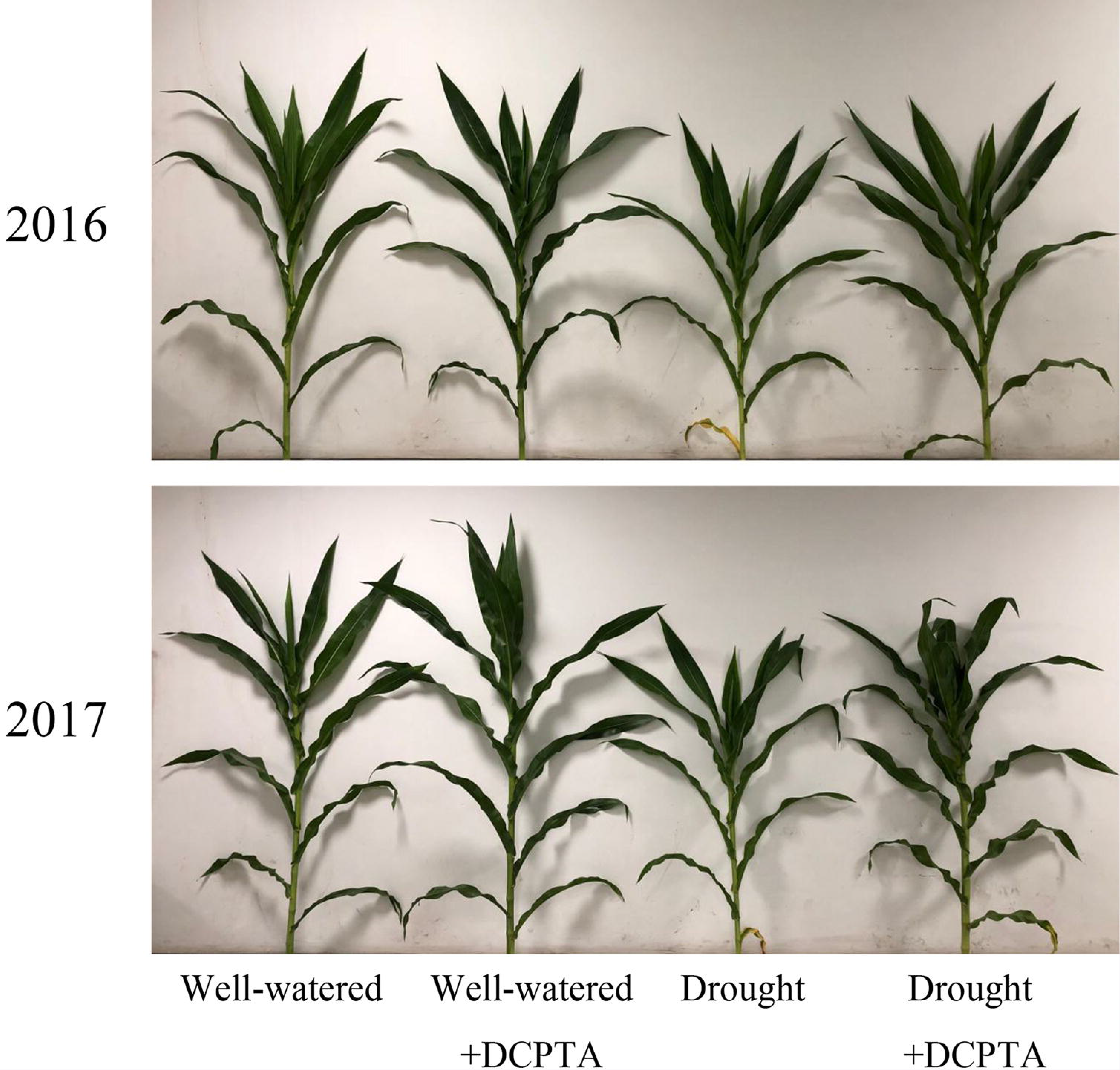
Leaf phenotypic features of the maize seedlings after 20 days of treatment with drought and/or DCPTA in 2016 and 2017.

**Fig. 7.**
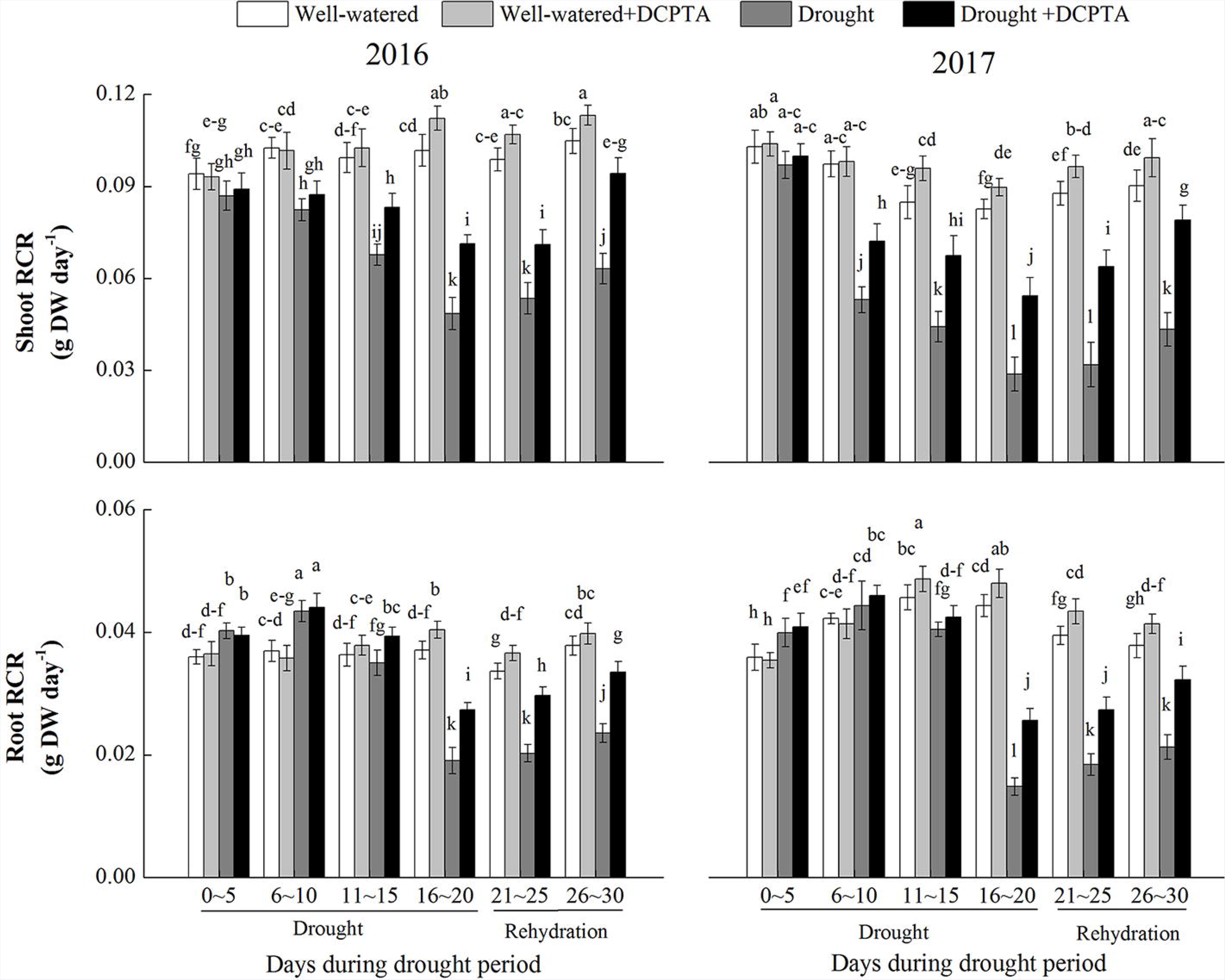
Effect of drought and/or DCPTA treatment on the relative growth rate (RGR) of the shoots and roots of the maize in 2016 and 2017. The data represent the means of independent measurements with five replicates, and the standard deviations are indicated by the vertical error bars. Values with the same letters on the bars are not significantly different at P<0.05 (LSD test).

### Root length density (RLD) and root surface area density (RSD)

Drought dramatically inhibited RLD and RSD in the 0–60 cm soil profile in 2016 and 2017 (Fig. 8). At day 20, there was a significant difference in RLD for the drought treatment and drought+DCPTA treatment in 20–40 cm in 2016 and in 20–50 cm in 2017, respectively. On day 30, the RLD declines were partially reversed by rehydration. On day 30, there was a significant difference in RLD for the drought treatment and drought+DCPTA treatment in 0–50 cm in both 2016 and 2017. On day 30, there was a significant (P > 0.05) difference in RLD for the well-watered treatment and well-watered+DCPTA treatment in 0–20 cm in both 2016 and 2017. On day 20, there was a significant difference in RLD between the drought treatment and drought+DCPTA treatment in 10–40 cm in 2016 and 10–50 cm in 2017. On day 30, the declines in RLD were partially reversed by rehydration. At day 30, there was a significant (P > 0.05) difference in RLD between the drought treatment and drought+DCPTA treatment in 0–40 cm in 2016 and 0–50 cm in 2017. On day 30, there was a significant difference between RLD for the well-watered treatment and well-watered+DCPTA treatment in 0–20 cm in 2016 and in 0–30 cm in 2017.

**Fig. 8.**
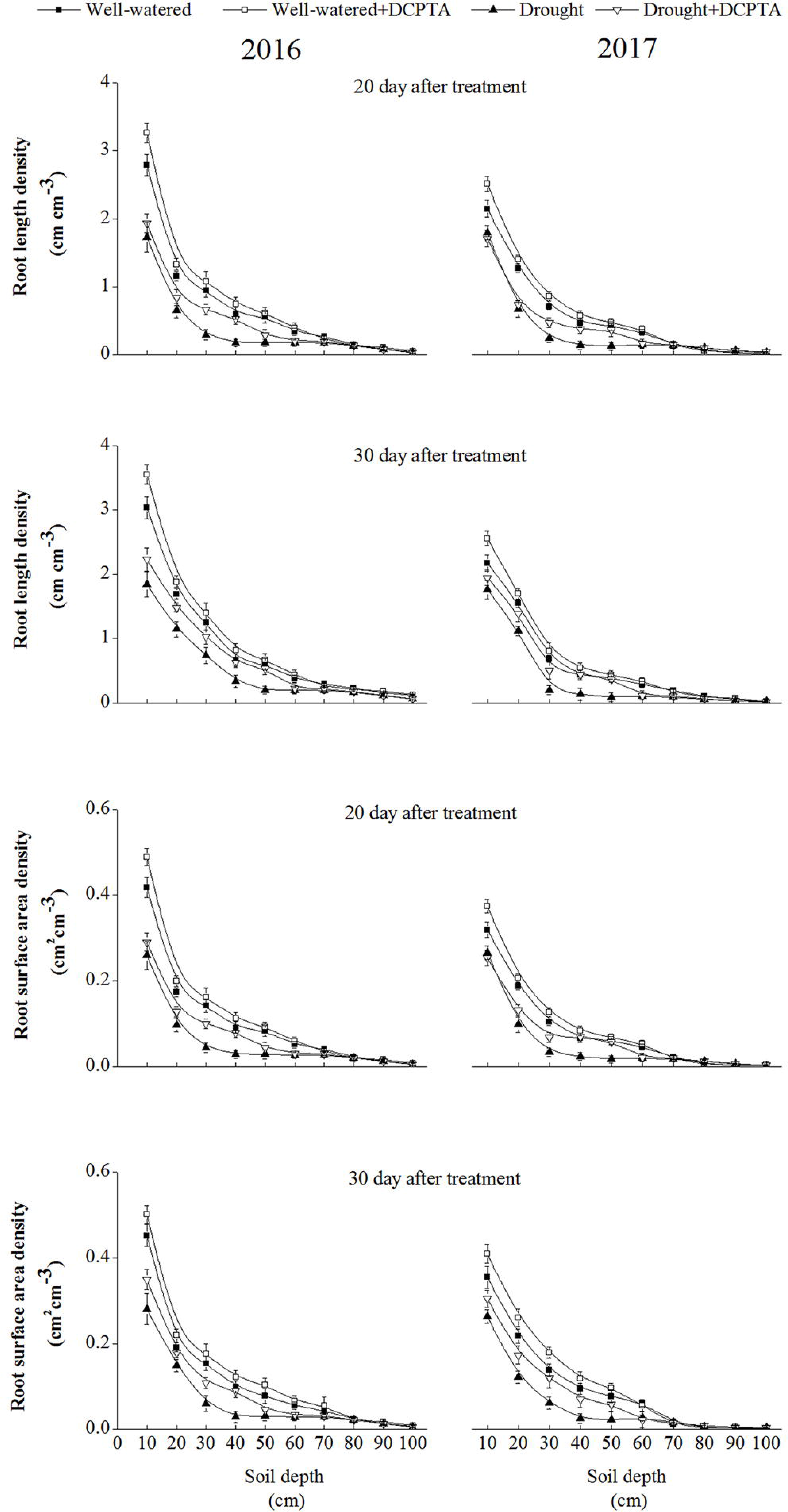
Effect of drought and/or DCPTA treatment on root length density (RLD) and root surface area density (RSD) for different soil depths on the 20^th^ day and 30^th^ day after treatment in 2016 and 2017.

### Root hydraulic conductivity, flow rate of root-bleeding sap and NO_3_^−^ delivery rates

The root hydraulic conductivity, flow rate of root-bleeding sap and NO_3_^−^ concentrations in the root-bleeding sap declined continuously during the drought period and recovered during rehydration (Fig. 9). In the drought treatment compared with the control, the root hydraulic conductivity, flow rate of root-bleeding sap and NO_3_^−^ delivery rates decreased by 34.21%, 75.69% and 76.58%, respectively, on day 20 and by 19.34%, 35.96% and 57.37%, respectively, on day 30 in 2016; these values decreased by 47.01%, 78.80% and 61.34%, respectively, on day 20 and by 29.03%, 33.79%, and 50.47%, respectively, on day 30 in 2017. However, the DCPTA application partially reversed the decline in root hydraulic conductivity, root-bleeding sap flow and NO_3_^−^ delivery rates caused by drought and resulted in a faster recovery after rehydration. In the drought+DCPTA treatment compared with the control, the root hydraulic conductivity, root-bleeding sap flow and NO_3_^−^ delivery rates decreased by 17.91%, 46.73% and 41.90%, respectively, on day 20 and by 9.87%, 18.80% and 23.51%, respectively, on day 30 in 2016; these values decreased by 26.60%, 47.66% and 37.06%, respectively, on day 20 and by 16.94%, 14.68% and 25.23%, respectively, on day 30.

**Fig. 9.**
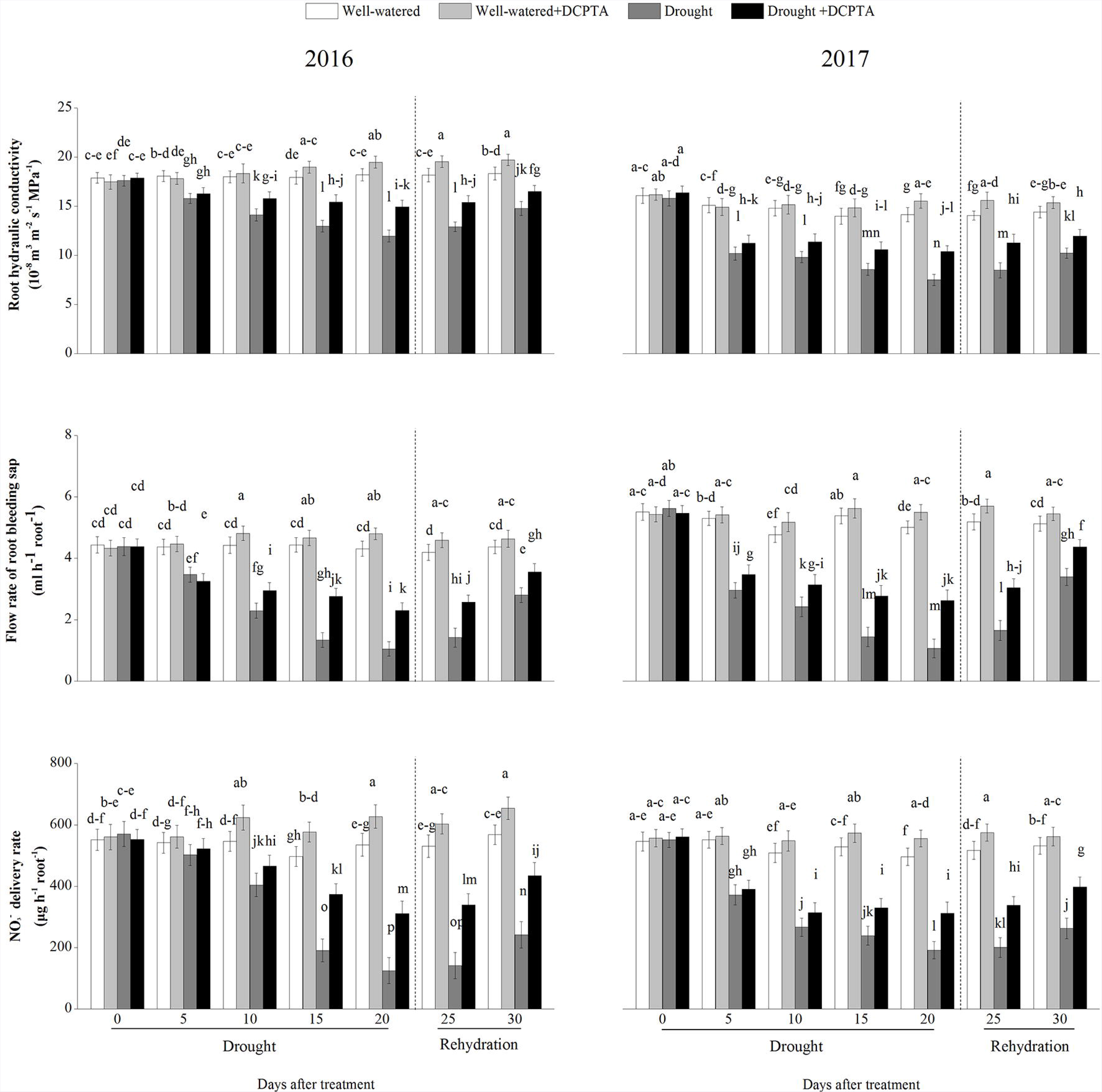
Effect of drought and/or DCPTA treatment on root hydraulic conductivity, the flow rate of root-bleeding sap and the NO_3_^−^ delivery rate in 2016 and 2017. The data represent the means of independent measurements with five replicates, and the standard deviations are indicated by the vertical error bars. Values with the same letters on the bars are not significantly different at P<0.05 (LSD test).

Under well-watered conditions, DCPTA significantly increased root hydraulic conductivity on the 15 ^th^, 20^th^, 25 ^th^ and 30^th^ days in 2016 and on the 20^th^ and 25 ^th^ days in 2017; DCPTA significantly increased the flow rate of root-bleeding sap on the 10^th^, 15^th^, 20^th^ and 25^th^ days in both 2016 and 2017; DCPTA significantly increased the NO_3_^−^ concentration in root-bleeding sap on the 10^th^, 15^th^, 20^th^, 25^th^ and 30^th^ days in 2016 and on the 10^th^, 15^th^, 20^th^ and 25^th^ days in 2017.

### Leaf water status (RLWC)

The RLWC declined continuously over the drought period and recovered during rehydration (Fig. 10). In the drought treatment compared with the control, RLWC decreased by 30.88% on day 20 and by 13.08% on day 30 in 2016 and decreased by 37.48% on day 20 and by 21.89% on day 30 in 2017. However, the DCPTA application partially reversed the decline in RLWC caused by drought and resulted in a faster recovery of the foliar RLWC contents after rehydration. In the drought+DCPTA treatment compared with the control, the RLWC decreased by 11.75% on day 20 and by 3.84% on day 30 in 2016 and decreased by 21.15% on day 20 and by 10.73% on day 30 in 2017. Under well-watered conditions, DCPTA application had no significant effect on the RLWC.

**Fig. 10.**
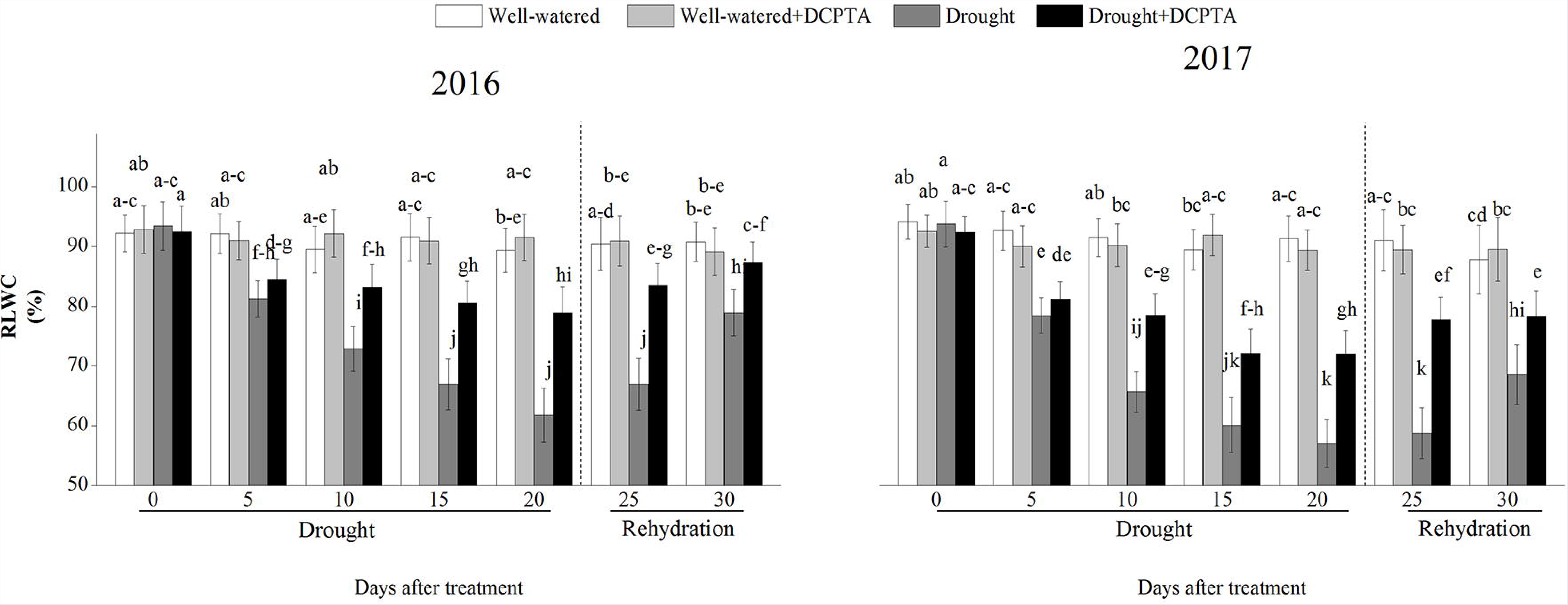
Effect of drought and/or DCPTA treatment on relative water content of the leaves (RLWC) in 2016 and 2017. The data represent the means of independent measurements with five replicates, and the standard deviations are indicated by the vertical error bars. Values with the same letters on the bars are not significantly different at P<0.05 (LSD test).

### Gas exchange parameters

Pn, Gs and Tr declined continuously over the drought period and recovered during rehydration (Fig. 11). In the drought treatment compared with the control, Pn, Gs and Tr decreased by 46.20%, 68.97% and 51.35%, respectively, on day 20 and by 35.06%, 46.30% and 38.17%, respectively, on day 30 in 2016 and decreased by 56.00%, 71.10% and 62.10%, respectively, on day 20 and by 45.89%, 44.02% and 43.79%, respectively, on day 30 in 2017. However, the DCPTA application partially reversed the decline in Pn, Gs and Tr caused by drought and resulted in a faster recovery of Pn, Gs and Tr rehydration. In the drought+DCPTA treatment compared with the control, Pn, Gs and Tr decreased by 23.41%, 37.39% and 20.95%, respectively, on day 20 and by 8.77%, 20.89% and 10.61%, respectively, on day 30 in 2016 and decreased by 35.31%, 32.03% and 34.11%, respectively, on day 20 and by 14.16%, 15.33% and 8.62%, respectively, on day 30. Under well-watered conditions, DCPTA significantly increased Pn on the 10^th^, 15^th^, 25^th^ and 30^th^ days in 2016 and on the 15^th^, 20^th^ and 25^th^ days in 2017. In addition, under well-watered conditions, DCPTA significantly increased Gs on the 15^th^, 20^th^, 25^th^ and 30^th^ days in 2016 and on the 10^th^, 15^th^, 20^th^, 25^th^ and 30^th^ days in 2017 and significantly increased Tr on the 10^th^, 15 ^th^, 20^th^, 25^th^ and 30^th^ days in both 2016 and 2017.

**Fig. 11.**
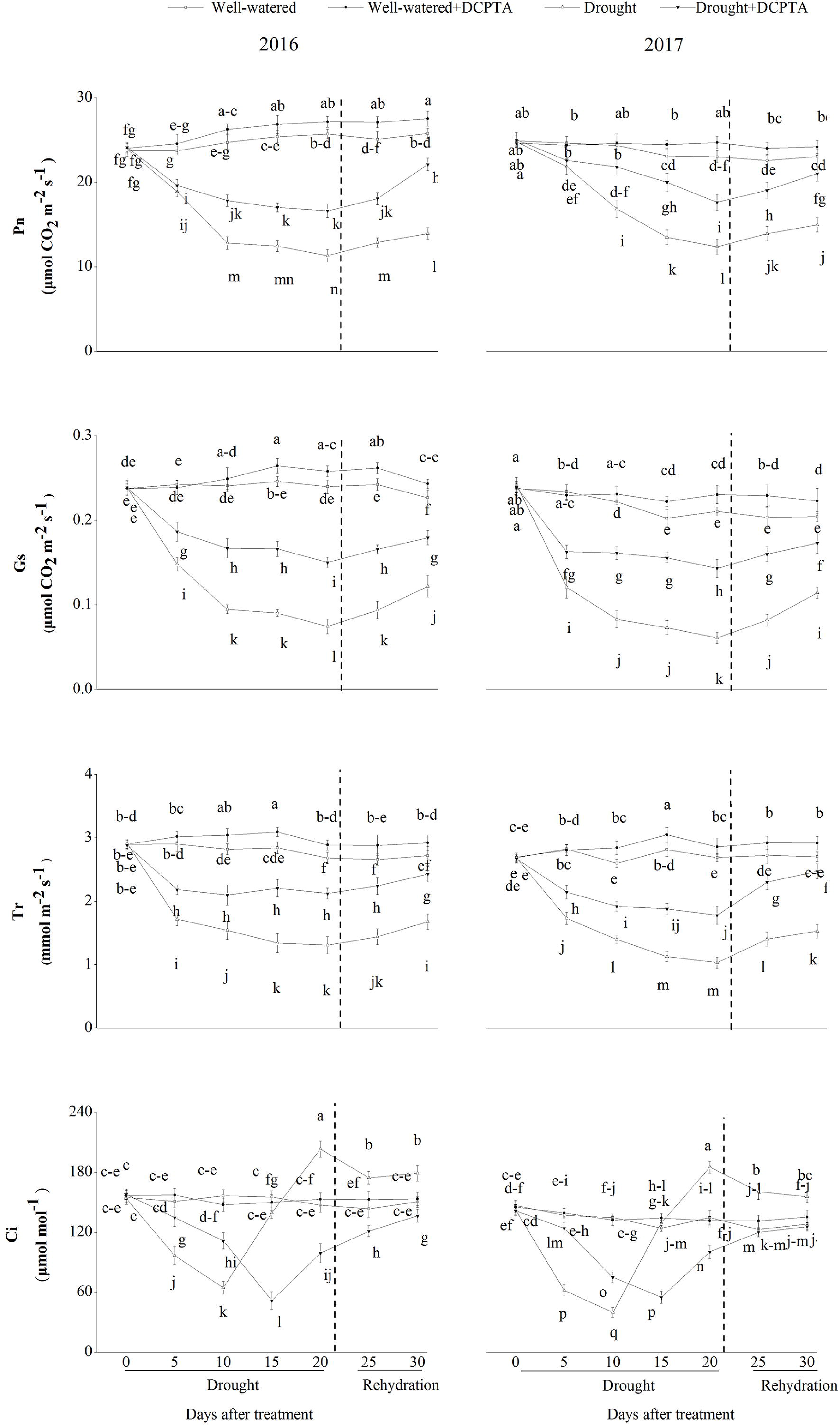
Effect of drought and/or DCPTA treatment on the gas exchange parameters in the maize leaves in 2016 and 2017. Pn, net photosynthetic rate; Gs, stomatal conductance; Tr, transpiration rate; Ci, intercellular CO_2_ concentration. The data represent the means of independent measurements with five replicates, and the standard deviations are indicated by the vertical error bars. Values with the same letters on the bars are not significantly different at P<0.05 (LSD test).

In 2016 and 2017, Ci showed the same variation tendency during the drought period. In the drought treatment, Ci declined over days 0-10, subsequently increased over days 10-20, and then decreased after rehydration. In the drought+DCPTA treatment, Ci declined over days 0-15 day and subsequently increased over days 15-30. Under well-watered conditions, DCPTA application had no significant effect on Ci.

### Chloroplast ultrastructure

Regardless of whether DCPTA was applied, the photosynthetic mesophyll cells of the non-stressed seedlings included a delimited cell wall containing chloroplasts. These chloroplasts had intact membranes and a regular arrangement of granal and stromal thylakoids, which were attached to the cell wall and exhibited typical ellipsoidal shapes (Fig. 12). However, in the stressed seedlings, the cell wall structure was incomplete and exhibited indistinct gradation, a lower density, and loose edges. Plasmolysis and degradation were also evident in part of the cell membrane. Moreover, the chloroplasts, which separated from the plasma membrane, were nearly round and swelled asymmetrically, the thylakoids were overly disorganized, and the thylakoid membranes were loose and showed an increased number of plastoglobules. In the PEG-6000+DCPTA treatment, the complete membrane structures of the chloroplasts were present, and the shapes of the chloroplasts changed slightly from elongated ellipses to ellipses close to the cell walls. A well-aligned internal lamellar system and fewer plastoglobules were observed in the leaves of the PEG-6000+DCPTA treatment compared with the leaves of the PEG-6000 treatment.

**Fig. 12.**
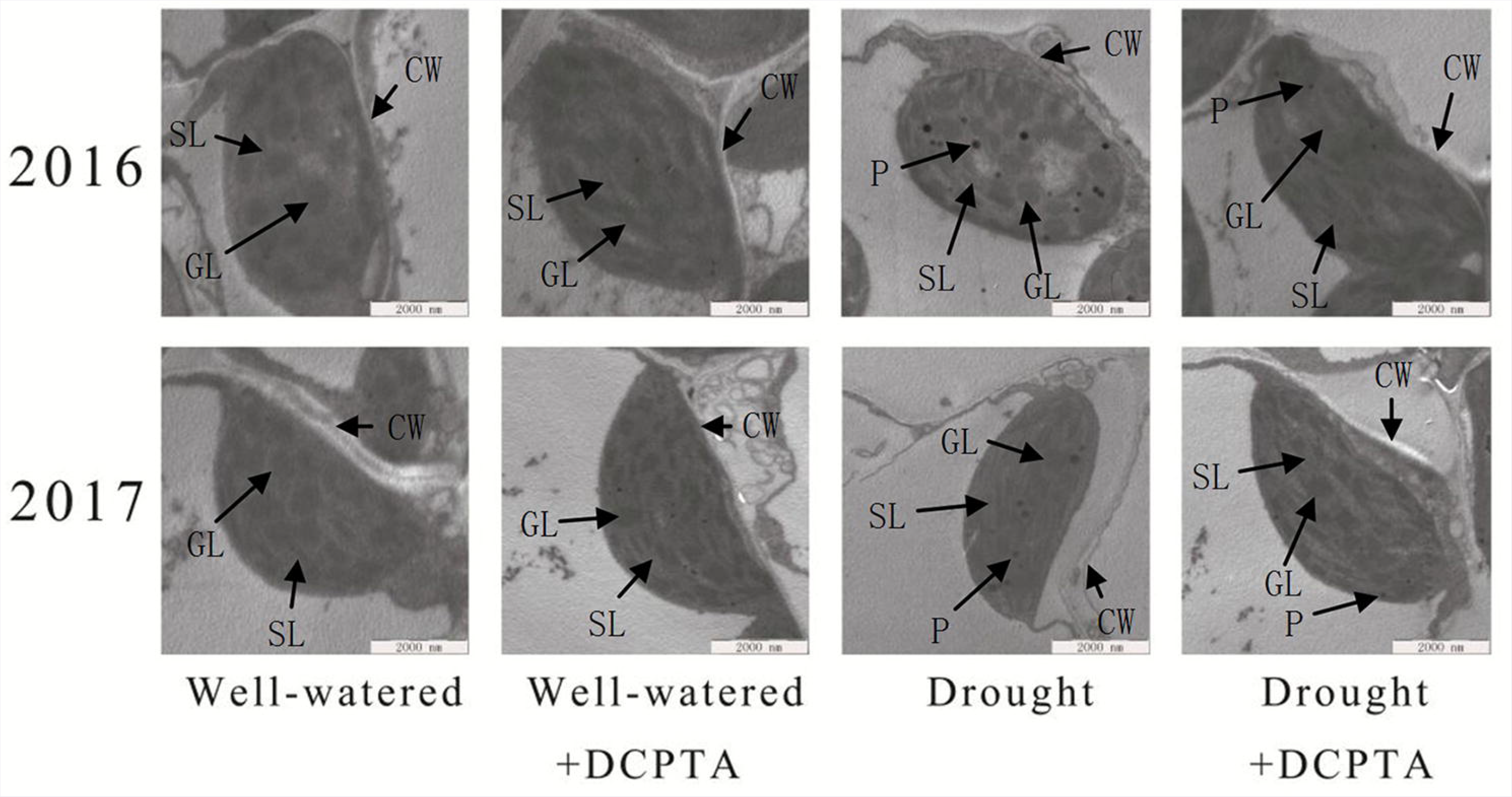
Ultrastructure of the photosynthetic apparatus in the maize leaves after 20 days of treatment with drought and/or DCPTA in 2016 and 2017. SL, stroma lamella; GL, grana lamellae; CW, cell wall; and P, plastoglobule. The scale bars for the photosynthetic apparatus represent 2000 nm.

### ICDH activity

ICDH activity declined continuously over the drought period and recovered during rehydration (Fig. 13). In the drought treatment compared with the control, ICDH activity decreased by 40.75% on day 20 and by 36.33% on day 30 in 2016 and decreased by 37.77% on day 20 and by 33.07% on day 30 in 2017. However, the DCPTA application partially reversed the decline in ICDH activity caused by drought and resulted in a faster recovery of ICDH activity after rehydration. In the drought+DCPTA treatment compared with the control, ICDH decreased by 24.92% on day 20 and by 19.94% on day 30 in 2016 and decreased by 19.88% on day 20 and by 13.63% on day 30 in 2017. Under well-watered conditions, the application of DCPTA significantly increased the foliar ICDH activity on the 10^th^, 15^th^, 20^th^ and 25^th^ days in 2016 and on the 15 ^th^, 20^th^, and 25 ^th^ days in 2017.

**Fig. 13.**
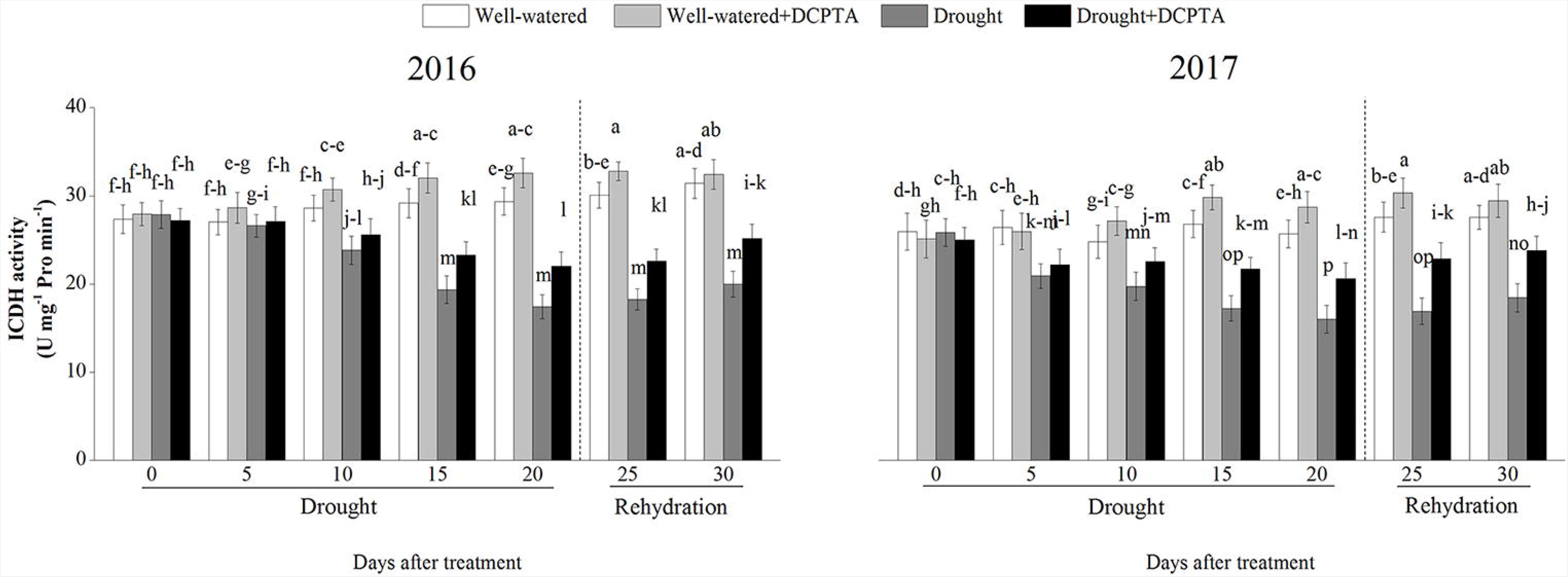
Effects of drought and/or DCPTA treatment on isocitrate dehydrogenase (ICDH) activity in 2016 and 2017. The data represent the means of independent measurements with five replicates, and the standard deviations are indicated by the vertical error bars. Values with the same letters on the bars are not significantly different at P<0.05 (LSD test).

### Contents of NO_3_^−^, NO_2_^−^ and NH_4_+

The foliar NO_3_^−^ and NO_2_^−^ contents declined continuously during the drought period and recovered during rehydration (Fig. 14). In the drought treatment compared with the control, the foliar NO_3_^−^ and NO_2_^−^ contents decreased by 39.82% and 38.27%, respectively, on day 20 and by 33.62% and 35.11%, respectively, on day 30 in 2016 and decreased by 57.97% and 32.27%, respectively, on day 20 and by 42.38% and 25.21%, respectively, on day 30 in 2017. However, the DCPTA application partially reversed the decline in the foliar NO_3_^−^ and NO_2_^−^ contents caused by drought and resulted in a faster recovery of the foliar NO_3_^−^ and NO_2_^−^ contents after rehydration. In the drought+DCPTA treatment compared with the control, the foliar NO_3_^−^ and NO_2_^−^ contents decreased by 25.94% and 23.31%, respectively, on day 20 and by 14.25% and 18.55%, respectively, on day 30 in 2016 and decreased by 33.86% and 18.26%, respectively, on day 20 and by 17.26% and 8.60%, respectively, on day 30 in 2017. In contrast, drought led to a marked elevation in the foliar NH_4_^+^ content. In the drought treatment compared with the control, the foliar NH_4_^+^ increased by 52.48% on day 20 and by 29.18% on day 30 in 2016 and increased by 98.68% on day 20 and by 72.21% on day 30 in 2017. In contrast, the DCPTA application suppressed the increase in the foliar NH_4_^+^ content induced by drought. In the drought+DCPTA treatment compared with the control, the foliar NH_4_^+^ increased by 20.83% on day 20 and by 13.33% on day 30 in 2016 and increased by 45.37% on day 20 and by 33.29% on day 30 in 2017.

**Fig. 14.**
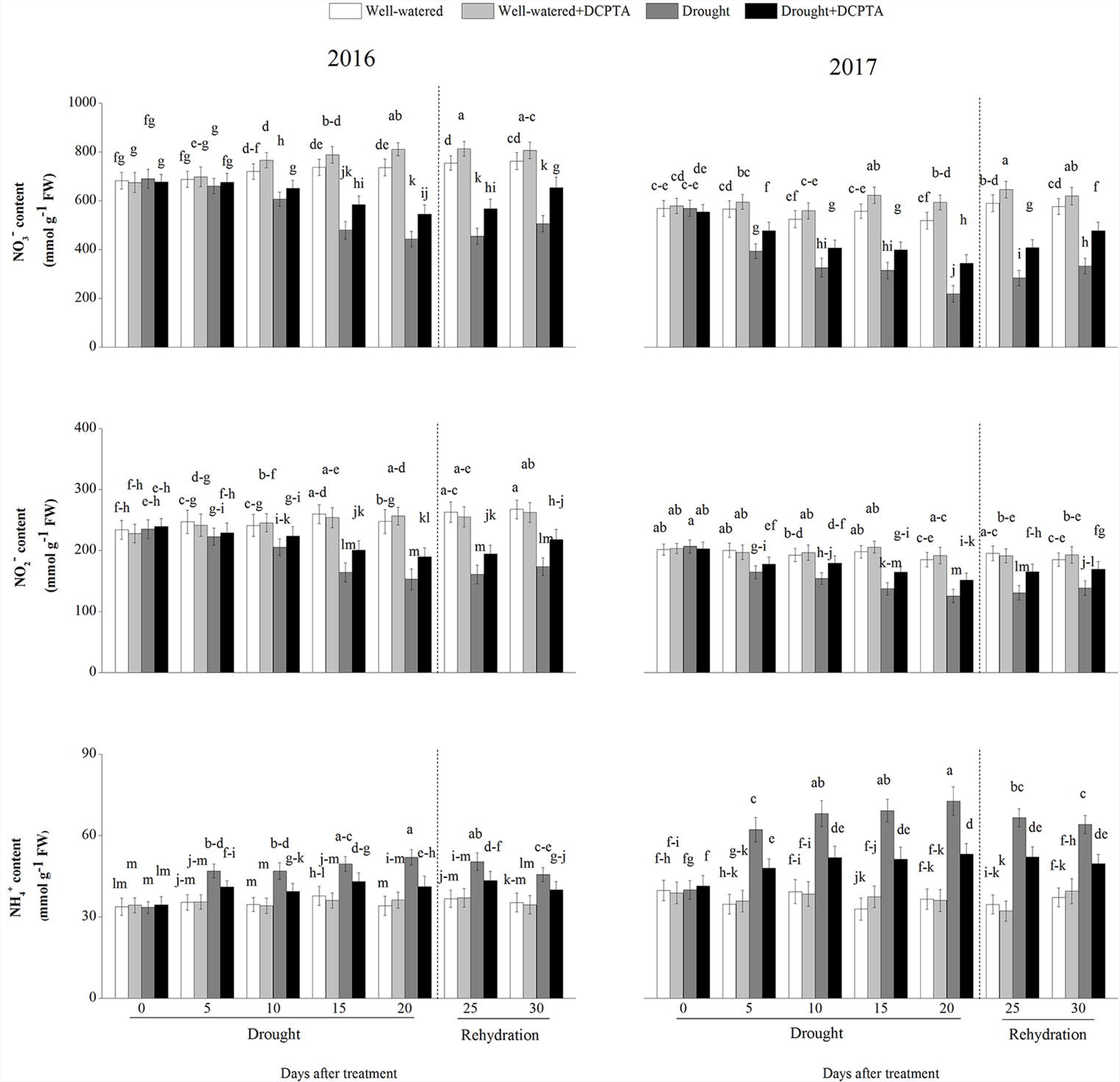
Effects of drought and/or DCPTA treatment on the nitrate (NO_3_^−^), nitrite (NO_2_^−^) and ammonium (NH_4_) contents in the leaves of maize in 2016 and 2017. The data represent the means of independent measurements with five replicates, and the standard deviations are indicated by the vertical error bars. Values with the same letters on the bars are not significantly different at P<0.05 (LSD test).

Under well-watered conditions, DCPTA significantly increased the foliar NO_3_^−^ content on the 20^th^ and 25 ^th^ days in 2016 and on the 15 ^th^, 20^th^, 25 ^th^ and 30^th^ days in 2017. However, the DCPTA application had no significant effect on the foliar contents of NO_2_^−^ and N¾^+^.

### Activities of NR and NiR

The activities of foliar NR and NiR declined continuously during the drought period and recovered during rehydration (Fig. 15). In the drought treatment compared with the control treatment, the activities of foliar NR and NiR decreased by 40.37% and 36.91%, respectively, on day 20 and by 34.42% and 29.82%, respectively, on day 30 in 2016 and decreased by 52.80% and 32.46%, respectively, on day 20 and by 37.36% and 21.28%, respectively, on day 30 in 2017. However, the DCPTA application partially reversed the declines in the activities of foliar NR and NiR caused by drought and resulted in a faster recovery of the foliar NR and NiR activities after rehydration. In the drought+DCPTA treatment compared with the control, the activities of foliar NR and NiR decreased by 15.54% and 14.21%, respectively, on day 20 and by 10.79% and 7.79%, respectively, on day 30 in 2016 and decreased by 25.10% and 12.39%, respectively, on day 20 and by 13.55% and 4.96%, respectively, on day 30 in 2017. Under well-watered conditions, DCPTA significantly increased the foliar NR activity on the 10^th^, 15 ^th^ and 25 ^th^ days in 2016 and on the 10^th^, 15 ^th^, 20^th^ and 25^th^ days in 2017. Similarly, under well-watered conditions, DCPTA significantly increased the foliar NiR activity on the 10^th^ and 25^th^ days in 2016 and on the 15^th^ and 20^th^ days in 2017.

**Fig. 15.**
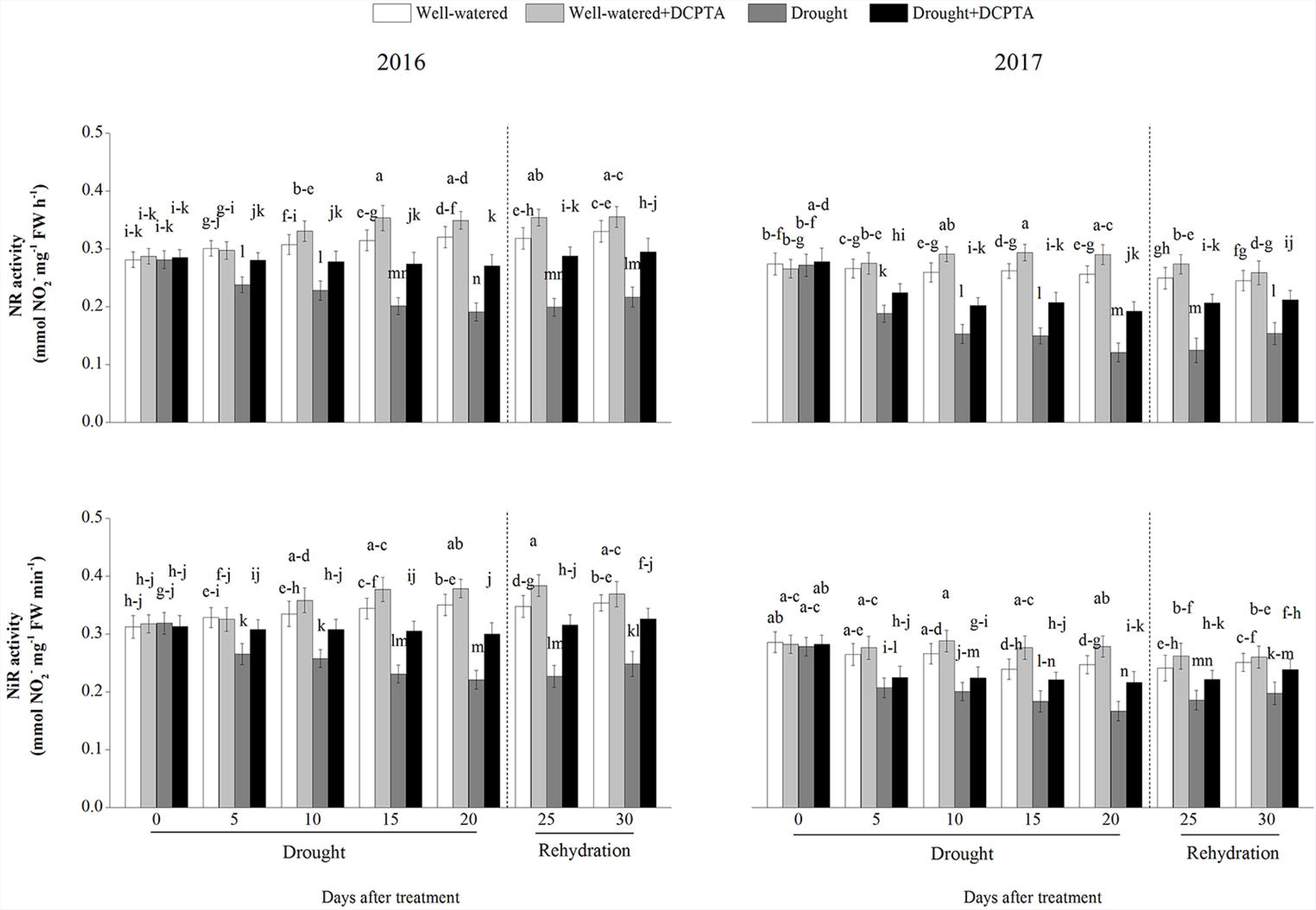
Effects of drought and/or DCPTA treatment on nitrate reductase (NR) and nitrite reductase (NiR) activities in the leaves of maize in 2016 and 2017. The data represent the means of independent measurements with five replicates, and the standard deviations are indicated by the vertical error bars. Values with the same letters on the bars are not significantly different at P<0.05 (LSD test).

## Activities of GS, GOGAT and GDH

The activities of foliar GS and GOGAT first increased and then decreased during the drought period and recovered during rehydration (Fig. 16). In the drought treatment compared with the control, the activities of foliar GS and GOGAT decreased by 40.69% and 60.62%, respectively, on day 20 and by 33.65% and 51.39%, respectively, on day 30 in 2016 and decreased by 47.57% and 66.84%, respectively, on day 20 and by 37.96% and 56.77%, respectively, on day 30 in 2017. However, the DCPTA application partially reversed the decline in the foliar GS and GOGAT activities caused by drought and resulted in a faster recovery of the foliar GS and GOGAT activities after rehydration. In the drought+DCPTA treatment compared with the control, the foliar GS and GOGAT activities decreased by 19.87% and 33.37%, respectively, on day 20 and by 10.99% and 15.17%, respectively, on day 30 in 2016 and decreased by 23.56% and 33.02%, respectively, on day 20 and by 14.93% and 18.20%, respectively, on day 30 in 2017. Likewise, the DCPTA application alone caused an increase in the foliar GS and GOGAT activities. In contrast, drought led to marked increases in the activities of foliar NAD-GDH and NADH-GDH. In the drought treatment compared with the control, the activities of foliar NAD-GDH and NADH-GDH increased by 87.16% and 150.92%, respectively, on day 20 and by 84.01% and 134.71%, respectively, on day 30 in 2016 and increased by 103.99% and 137.36%, respectively, on day 20 and by 96.13% and 111.20%, respectively, on day 30 in 2017. However, the DCPTA application partially reversed the increases in the activities of foliar NAD-GDH and NADH-GDH caused by drought and resulted in a faster recovery of the foliar NAD-GDH and NADH-GDH activities after rehydration. In the drought+DCPTA treatment compared with the control, the activities of foliar NAD-GDH and NADH-GDH increased by 49.55% and 92.59%, respectively, on day 20 and by 36.52% and 49.51%, respectively, on day 30 in 2016 and increased by 46.90% and 80.94%, respectively, on day 20 and by 35.43% and 45.60%, respectively, on day 30 in 2017.

**Fig. 16.**
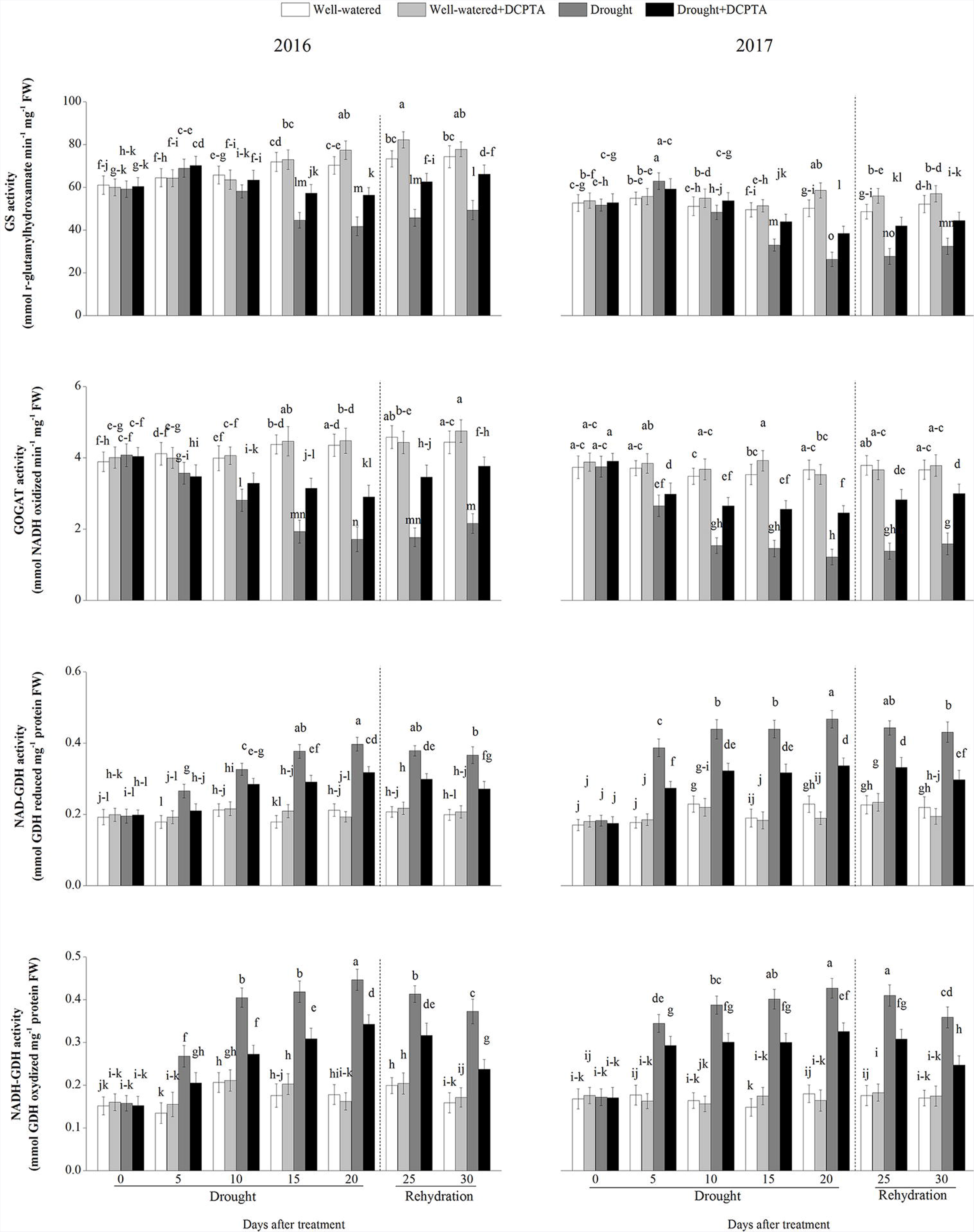
Effects of drought and/or DCPTA treatment on the glutamine synthetase (GS), glutamate synthase (GOGAT) and glutamate dehydrogenase (GDH) activities in the leaves of maize in 2016 and 2017. The data represent the means of independent measurements with five replicates, and the standard deviations are indicated by the vertical error bars. Values with the same letters on the bars are not significantly different at P<0.05 (LSD test).

Under well-watered conditions, DCPTA significantly increased the foliar GS activity on the 20^th^ and 25 ^th^ days in 2016 and on the 10^th^ 20^th^ and 25 ^th^ days in 2017. Similarly, under well-watered conditions, DCPTA significantly increased the foliar GOGAT activity on day 15 in 2017 and significantly increased the foliar NAD-GDH activity on day 15 in 2016

### Activities of AlaAT and AspAT

The activities of foliar AlaAT and AspAT first increased and then continuously decreased during the drought period and recovered during rehydration (Fig. 17). In the drought treatment compared with the control, the activities of foliar AlaAT and AspAT decreased by 44.18% and 65.51%, respectively, on day 20 and by 36.43% and 38.52%, respectively, on day 30 in 2016 and decreased by 56.80% and 54.92%, respectively, on day 20 and by 41.43% and 41.29%, respectively, on day 30 in 2017. However, the DCPTA application partially reversed the decline in the activities of foliar AlaAT and AspAT caused by drought and resulted in a faster recovery of the foliar AlaAT and AspAT activities after rehydration. In the drought+DCPTA treatment compared with the control, the activities of foliar AlaAT and AspAT decreased by 17.45% and 35.39%, respectively, on day 20 and by 15.84% and 17.17%, respectively, on day 30 in 2016 and decreased by 32.49% and 37.82%, respectively, on day 20 and by 17.46% and 18.21%, respectively, on day 30 in 2017.

**Fig. 17.**
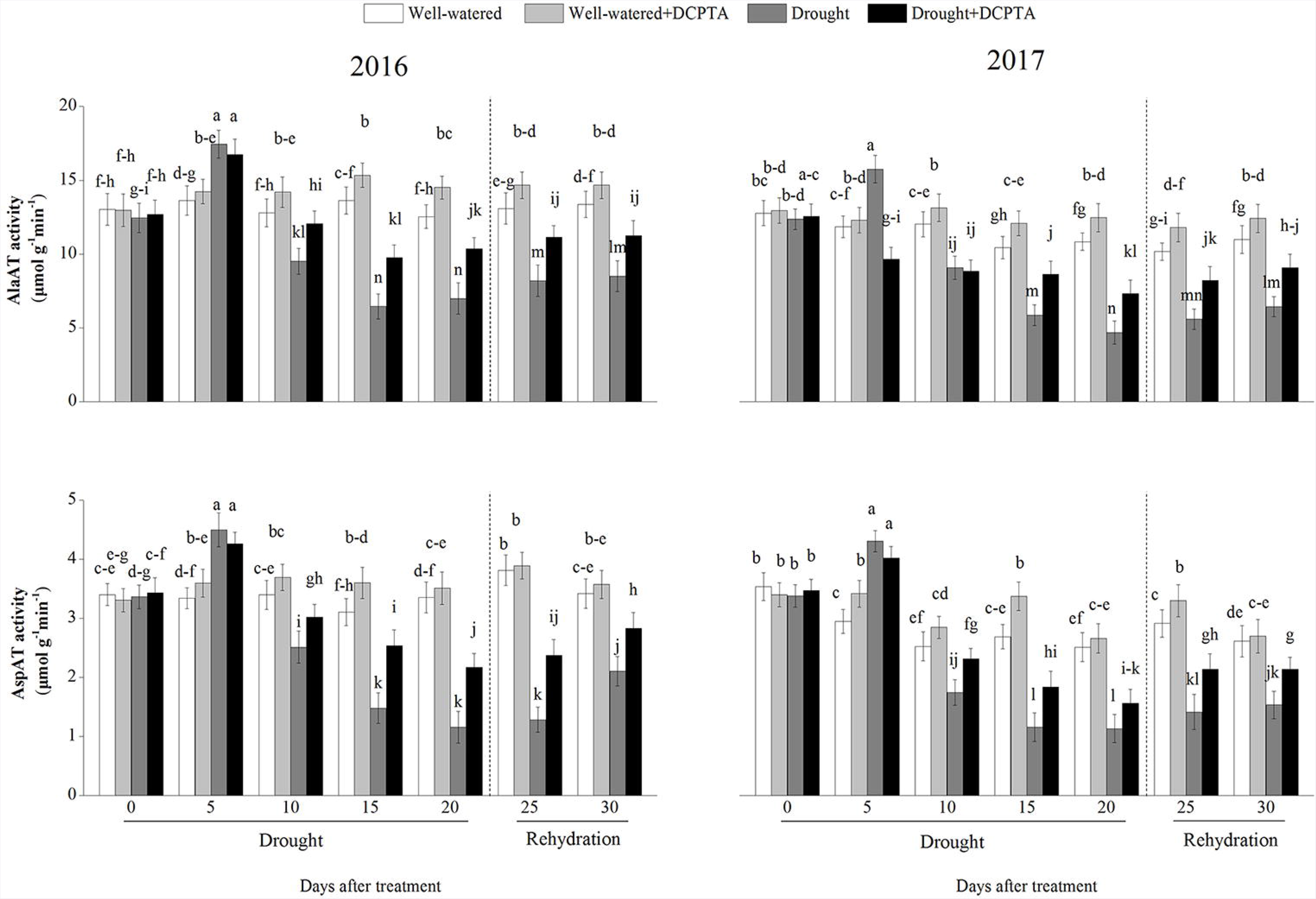
Effects of drought and/or DCPTA treatment on the alanine aminotransferase (AlaAT) and aspartate aminotransferase (AspAT) activities in the leaves of maize in 2016 and 2017. The data represent the means of independent measurements with five replicates, and the standard deviations are indicated by the vertical error bars. Values with the same letters on the bars are not significantly different at P<0.05 (LSD test). Fig. 18 Effects of drought and/or DCPTA treatment on the protease activity and protein and free amino acid contents in the leaves of maize in 2016 and 2017. The data represent the means of independent measurements with five replicates, and the standard deviations are indicated by the vertical error bars. Values with the same letters on the bars are not significantly different at P<0.05 (LSD test).

Under well-watered conditions, DCPTA significantly increased the foliar AlaAT activity on the 10^th^, 15^th^, and 20^th^ days in 2016 and on the 10^th^, 15^th^, 20^th^ and 25^th^ days in 2017. Similarly, under well-watered conditions, DCPTA significantly increased the foliar AspAT activity on day 15 in 2016 and on the 5^th^, 10^th^, 15^th^ and 25^th^ days in 2017.

### Protease activity and contents of proteins and free amino acids

The protease activity and free amino acid contents increased continuously during the drought period and decreased during rehydration (Fig. 18). In the drought treatment compared with the control, the protease activity and free amino acid contents increased by 122.48% and 88.92%, respectively, on day 20 and by 55.03% and 34.51%, respectively, on day 30 in 2016 and increased by 145.15% and 78.58%, respectively, on day 20 and by 57.08% and 43.29%, respectively, on day 30 in 2017. However, the DCPTA application partially reversed the increases in the protease activity and free amino acid contents caused by drought.

**Fig. 18.**
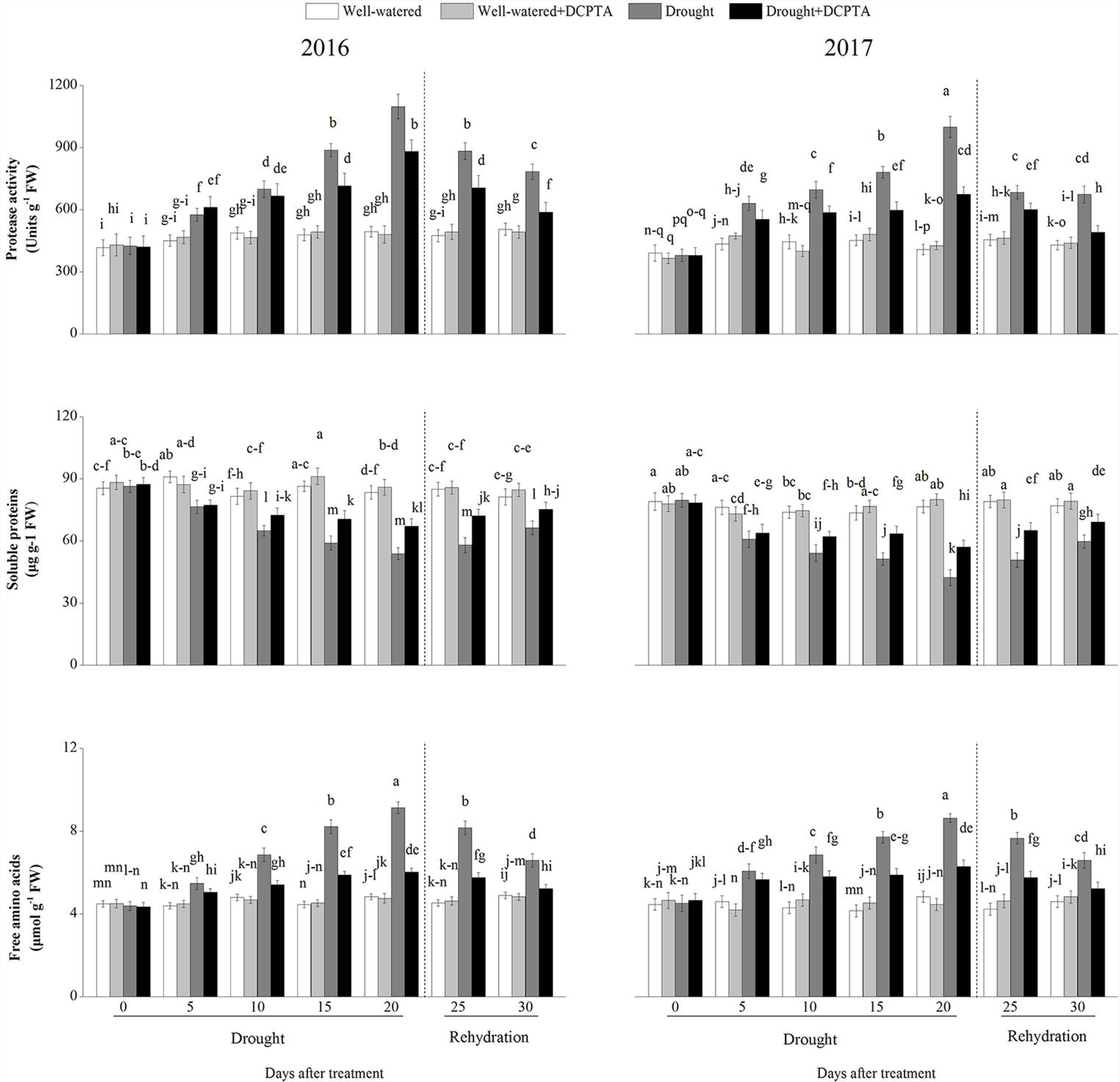
Effects of drought and/or DCPTA treatment on the protease activity and protein and free amino acid contents in the leaves of maize in 2016 and 2017. The data represent the means of independent measurements with five replicates, and the standard deviations are indicated by the vertical error bars. Values with the same letters on the bars are not significantly different at P<0.05 (LSD test).

In the drought+DCPTA treatment compared with the control, the protease activity and free amino acid contents increased by 78.47% and 24.77%, respectively, on day 20 and by 16.46% and 6.62%, respectively, on day 30 in 2016 and increased by 65.41% and 30.24%, respectively, on day 20 and 14.50% and 13.57%, respectively, on day 30 in 2017.

In contrast, drought led to a marked decrease in the foliar protein content. In the drought treatment compared with the control, the foliar protein content decreased by 35.51% on day 20 and by 18.32% on day 30 in 2016 and decreased by 44.81% on day 20 and by 22.50% on day 30 in 2017. In contrast, the DCPTA application suppressed the increase in the foliar protein content induced by drought. In the drought+DCPTA treatment compared with the control, the foliar protein content increased by 19.54% on day 20 and by 7.25% on day 30 in 2016 and increased by 25.46% on day 20 and by 10.30% on day 30 in 2017.

Significant differences between the foliar protein contents of the well-watered treatment and well-watered+DCPTA treatment were observed on day 10 in 2017 and on day 15 in 2017, and significant differences between the free amino acid contents of these treatments were observed on day 10 in 2017.

## Discussion

One of the most clear and consistent effects of drought on crops is the inhibition of growth and yield (Sinclair, 2011). In this experiment, exogenous DCPTA partly mitigated the reductions in plant growth and yield induced by drought, as expressed by the stable soot and root RGR, GN and GY (Fig. 5, Fig. 6 and Fig. 7). Moreover, exogenous DCPTA promoted growth and yield under well-watered conditions.

N is a necessary macro-nutrient element for plants and a main limiting factor in plant growth and development (Kusano *et al.,* 2011). The pre-female inflorescence emergence stage is the important stage determining maize yield (Talaat *et al.,* 2015). The inhibition of maize growth induced by drought could be partly attributed to nitrogen metabolism during the pre-female inflorescence emergence stage. NO_3_^−^ is the main nitrogen source assimilated by higher plants in agricultural soils (Robredo *et al.,* 2011). Similar to previous reports for tomato (Sánchez-Rodríguez *et al.,* 2011) and wheat (Fresneau *et al.,* 2007), drought significantly diminished the NO_3__ content in maize leaves in both the DCPTA-treated and non-treated leaves (Fig. 14). This decrease may be explained by drought-induced inhibitions in nitrate uptake from the roots and/or nitrate transport. However, the reduction in the non-treated leaves was greater than that in the DCPTA-treated leaves in this study.

Maintaining the supply of nutrients and water from the soil to crop depends on root morphology, which can be characterized by RLD and RSD (Kamran *et al.,* 2018). Previous studies have reported that DCPTA promotes root development in tomatoes (Keithly *et al,* 1990a). In this study, there was a significant difference between the RLD and RSD of the well-watered treatment and DCPTA treatment in 0-20 cm and 0-30 cm in 2016 and 2017 on day 30 (Fig. 8). This result suggests that DCPTA promoted root development in maize under well-watered conditions. Interestingly, under drought conditions, the DCPTA application significantly increased RLD, RSD and the root RGR. These results indicate that the DCPTA application also promoted maize root growth and improved the spatial and temporal distribution of roots, which was beneficial to NO_3_^−^ uptake under drought conditions.

The xylem sap transports water and nutrients from the roots throughout the plant and depends on transpiration intensity and root pressure. The increased root hydraulic conductivity and flow rate of root-bleeding sap induced by the DCPTA application may, due to the enhanced root pressure, which depend on physiological activity of the whole root system (Fig. 9) (Noguchi *et al.,* 2005; Lui *et al,* 2014). In addition, the NO_3__ delivery rate in the presence of DCPTA was significantly higher than that without DCPTA under drought conditions, which may partly result from the improved NO_3__ absorption and enhanced root pressure induced by DCPTA. The stable RLWC in the drought+DCPTA treatment suggests an abundant supply water to the aboveground parts, and balanced transpirational loss and water uptake under drought conditions; as a result, the DCPTA-treated plants maintained Gs, which reduced the leaf epidermal resistance and promoted the mass flow of water to the leaf surface and the transportation of the NO3_ required for N metabolism in leaves (Fig. 10 and Fig. 11). Under drought conditions, increases in the foliar NO3_ were observed in the DCPTA-treated plants (Fig. 14).

Whether stomatal or non-stomatal factors are the main cause of a reduced Pn may be determined by changes in Gs and Ci (Bethke and Drew, 1992). During the early period of drought, the change of Ci were accompanied by continuously declined Gs, then Gs decreases but Ci shows an increase (Fig. 11).

Thus, the decrease of the Pn in drought-treated plants was mainly attributed to stomatal limitations firstly, and then, non-stomatal limitations induced by the damage of photochemical mechanism, partly reflected by damaged chloroplast (Fig. 12). However, DCPTA application maintains relatively high Gs, ensuring the availability of CO_2_ for the carbon reduction cycle. Simultaneously, the DCPTA application delayed the increase in *Ci* and protected the chloroplast ultrastructure against drought-induced oxidative damage, which suggests that DCPTA can protect the photochemical mechanism and, as a result, ensures a more efficient photosynthesis process after rehydration. Moreover, similar to previous studies on spruce (Keithly *et al.,* 1990a), sugar beets (Keithly *et al,* 1990b) and guayule (Wan and Mendoza, 1992), DCPTA application also promoted photosynthesis under well-watered conditions.

In most plants, nitrate reduction occurs in leaves. NO3, after being taken up into the leaf cell, is converted to NH_4_^+^ by two successive steps catalysed by NR and NiR. NR, the rate-limiting enzyme of nitrogen assimilation, is highly sensitive to stress (Plett *et al.,* 2016). Similar to previous studies on wheat and barley, the NR activity continuously declined in response to drought (Fig. 15) (Robredo *et al.,* 2011; Fresneau *et al.,* 2007). As a typical nitrate-induced enzyme, NR activity is primarily regulated by the NO_3_^−^ concentration in the leaves (Chamizo-Ampudia *et al.,* 2017). The up-regulation of foliar NR activity in the drought+DCPTA treatment may result from the increase in the foliar NO_3_^−^ content (Fig. 14). Moreover, the reduction in the foliar NiR activity under drought conditions was significantly reversed by the application of DCPTA, which may be because the DCPTA-stabilized photosynthesis resulted in a sufficient supply of Fdred (Fig. 11 and Fig. 15), thus promoting the conversion of NO_2_^−^ to N¾+. The present results indicate that DCPTA pretreatment could maintain a high NO_3_^−^ assimilation ability in maize under drought conditions.

Although the foliar NR and NiR activities declined during the drought period, the foliar NH_4_^+^ content exhibited an increasing tendency in our experiment (Fig. 14). This increase may be associated with the glycine oxidation in activated photorespiration, which is induced by decreases in *Ci* levels under drought conditions (Wang *et al.,* 2012). The increased *Gs* induced by DCPTA was beneficial to the increase in CO_2_ in the cellular spaces of the leaf, implying that photorespiration was partly alleviated (Fig. 11).

In plants cells, excessive levels of NH_4_+ are destructive, and the major NH_4_+ assimilation pathway is the GS/GOGAT cycle in higher plants. When the GS/GOGAT cycle is suppressed and the NH_4_^+^ content rises continuously under stress, NH_4_^+^ could serve as a substrate to form glutamate via the reversible amination of 2-OG by GDH catalyse, although the enzyme has a lower affinity for NH_4_^+^ (Fontaine *et al.,* 2012). In general, drought inhibited N¾^+^ assimilation (Naya *et al.,* 2007). During the early period of drought, GDH activity increased sharply, GS activity increased slightly, and GOGAT remained stable (Fig. 16). These results suggest that accelerated NH_4_^+^ assimilation in maize may be an adaptive mechanism to produce more glutamate and eliminate the accumulation of excess foliar NH_4_^+^. Subsequently, the GS and GOGAT activities decreased, which may have resulted from an inadequate supply of energy and 2-OG because of photosynthetic inhibition and decreased ICDH activity (Fig. 13). GDH activity decreased with drought, which may be due to the oxidative degradation of GS caused by the overproduction of reactive oxygen species (ROS) induced by drought conditions (Xia *et al.,* 2015).

The DCPTA application altered the major N¾^+^ assimilation pathway, maintained the GOGAT/GS cycle and suppressed the GDH pathway, which may have contributed to maintaining the conversion of NH_4_^+^ to glutamine and the subsequent formation of glutamate from glutamine. This result may occur because the photosynthetic stability and ICDH activity induced by the DCPTA application promoted 2-OG synthesis and the reducing power (i.e., NADPH, ATP, or Fd_re_d) in plants during the drought period, thus providing the GS/GOGAT cycle with relatively sufficient substrates and energy and favouring the enhancement of foliar GOGAT and GS activities (Du *et al.,* 2016). As a result, with the application of DCPTA, drought had less of an effect on the activities of GS and GOGAT.

Although DCPTA promoted NO_3_^−^ assimilation, as expressed by the increased NR and NiR activities (Fig. 15), this treatment compared to the drought treatment caused significant decreases in the NH_4_^+^ content, which means that exogenous DCPTA resulted in the integration of NH_4_^+^ into the structure of organic compounds, thereby contributing to the reduction in the NH_4_^+^ content. Therefore, the DCPTA application effectively modulated the activities of ICDH, GS, GOGAT and GDH and accelerated the conversion of NH_4_+ to glutamate, which is the precursor of other amino acids.

Transamination is a key step in the biosynthesis of various amino acids from glutamate, with the availability of C skeletons from the Krebs cycle (Hodges, 2002). In our studies, both the aminotransferases studied, AlaAT and AspAT, showed increased activity in maize during the early drought period (Fig. 17). Such increases in aminotransferases activities under drought conditions might help in the synthesis of increased amounts of amino acids that act as compatible cytoplasmic solutes and protect cell organelles and biomolecules, thus reducing the adverse effects of drought on maize (Munns and Tester, 2008). Subsequently, the AlaAT and AspAT activities decreased, which may be attributable to the weakened GS/NADH-GOGAT pathway (Fig. 16) (Gangwar and Singh, 2011). Moreover, stable aminotransferase activities were observed in DCPTA-treated plants. This finding may be associated with increased GS/GOGAT activities, which can generate more glutamate to serve as a substrate for transamination reactions in maize treated with DCPTA under drought conditions.

Most soluble proteins are enzymes that participate in various metabolic pathways in plants; therefore, the soluble protein content is considered one of the most important indices reflecting the overall metabolic level in plants. Protein synthesis in plants is very sensitive to abiotic stresses and is positively correlated with stress tolerance (Fresneau *et al.,* 2007). Free amino acids are the building blocks of proteins. Drought increased the free amino acid contents, which may mainly be attributed to the increased AlaAT and AspAT activities in the early drought period and the subsequent promotion of protein degradation (Fig. 18) (Yang *et al.,* 2013). However, DCPTA-treated seedlings maintained higher soluble protein levels and lower free amino acid levels than did non-DCPTA-treated seedlings in response to drought. This result may occur because DCPTA inhibited protein degradation by stable protease activities and maintained protein stability, ensuring the series of physiological and biochemical processes that occur normally under stress conditions. Additionally, the DCPTA application increased the amino acid contents under well-watered conditions, which may be attributable to the promoted biosynthesis and accumulation of amino acids, which ultimately improved plant growth and development (Talaat and Shawky, 2016).

## Conclusions

The present study suggested that DCPTA treatment increased NO3_ uptake and the long-distance transportation of NO3^_^ from the roots to the leaves via the production of excess roots and maintained a stabilized transpiration rate. The increased foliar NO3^_^ content up-regulated NR activity and maintained a high N assimilation ability that was restrained by drought. Exogenous DCPTA effectively regulated the ICDH, GS, GOGAT and GDH activities to speed up the conversion of NH_4_^+^ to Glu, reduced the toxicity of excess NH_4_^+^ to the plant, and accelerated the synthesis of proteins and amino acids. Moreover, DCPTA treatment maintained increased the photosynthetic capacity, supply nitrogen metabolism of energy and carbon skeleton thus alleviating the inhibition of growth by drought in maize.

## Acknowledgements

This work was supported by the National Key Research and Development Program of China (2016YFD0300103), the National Key Research and Development Program of China (2017YFD0300506), and the “Academic Backbone” Project of Northeast Agricultural University (17XG23).

